# Identification of Human Gut Microbiome–Derived Peptides Targeting Biofilm-Specific Lectin Proteins of *Pseudomonas aeruginosa*

**DOI:** 10.64898/2026.02.10.704983

**Authors:** Ayush Amod, Ananya Anurag Anand, Sreyanki Chandra, Sarfraz Anwar, Mubashra, Sameer Srivastava, Sintu Kumar Samanta

## Abstract

The carbohydrate-binding proteins (LecA and LecB) present within the extracellular polymeric substance (EPS) matrix of *Pseudomonas aeruginosa* play an essential role in maintaining the structural integrity of the biofilms through interactions with the EPS polysaccharides. Therefore, targeting the above lectins can turn out to be one of the promising strategies for disrupting *P. aeruginosa* biofilms. In the current study, we investigated the potency of antimicrobial peptides (AMPs) produced by the human gut microbiome in targeting LecA and LecB proteins of *P. aeruginosa*. Initially, a comprehensive *in-silico* pipeline was developed to identify and characterize putative antibacterial and antibiofilm AMPs produced by the human gut microbiome. These AMPs were then subsequently studied for their interaction with the lectin proteins through molecular docking, MM-GBSA, residue analysis, and molecular dynamics (MD) simulation. Among the studied peptides, amp21 and amp24 exhibited the strongest interactions with the lectin protein, occupying binding sites overlapping with key active-site residues previously reported for raffinose binding. amp6, amp21, and amp24 were selected for *in vitro* validation based on the MD simulation results of both LecA and LecB proteins. The above selected peptides exhibited minimal hemolytic activity across the tested concentration range. amp21 and amp6 were non-toxic to mammalian cells while amp24 demonstrated cytotoxicity only at higher doses. amp21 was found to be the most potent AMP and inhibited the growth of *P. aeruginosa* by ∼60% at 50 µg mL□¹. amp6 and amp21 resulted in a significant disruption of *P. aeruginosa* biofilms. Membrane permeabilization assays and scanning electron microscopy revealed that amp6, amp21, and amp24 damaged the bacterial cell membranes apart from compromising the integrity of the biofilm EPS matrix. Lastly, through *in-silico* studies, we designed ultrashort peptides (USPs) from the lead AMPs. The USPs (amp21.4 and amp24.2) exhibit superior antibiofilm efficacy compared to their parent AMPs. These findings establish human-gut microbiome-derived AMPs as promising candidates to target *P. aeruginosa* biofilms via inhibition of lectin proteins.

## 1. Introduction

Due to strong selection pressure driven by increasing antibiotic use and misuse over the last few years, a large number of bacterial species have evolved to acquire antimicrobial resistance (AMR) to a wide repertoire of antibiotics [Salam et al., 2023]. The World Health Organization (WHO) has identified AMR as one of the top ten global public health concerns, with recent estimates projecting that it can account for nearly 1.91 million deaths annually by 2050 [WHO 2020; Naghavi et al., 2024]. Bacterial biofilms substantially contribute to AMR by impeding antibiotic penetration, facilitating horizontal gene transfer among resident bacteria, and fostering metabolic heterogeneity. As a result, bacterial biofilms show increased tolerance to antibiotic agents, making them much harder to eradicate as compared to free-living planktonic cells [Hall-Stoodley et al., 2004]. Several studies report that the innate ability of ESKAPE pathogens to form biofilms results in broad-spectrum tolerance against conventional antibiotics, rendering current antimicrobial strategies almost ineffective [Sarkar et al., 2024].

Most antibacterial agents, including antibiotics, struggle to penetrate the biofilm’s EPS matrix, making biofilm cells 10–1000 times more tolerant to antimicrobials than their planktonic counterparts [Shree et al., 2023]. EPS matrix serves as a reservoir for several antibiotic-chelating enzymes, rendering the antibiotics non-functional. Moreover, the EPS matrix limits the penetration of antibiotics into the deeper layers of the biofilm [Singh and Amod et al., 2022]. Further, it promotes adhesion to the surface, cell-to-cell adhesion, and aggregation during biofilm formation [Flemming and Wingender, 2010].

The biofilm EPS matrix constitutes nearly 90% of total biofilm mass and is mainly comprised of extracellular DNA (eDNA), proteins, and polysaccharides [Fulaz et al., 2019]. One of the novel strategies to disrupt bacterial biofilms could be targeting structural proteins. Structural proteins, along with polysaccharides, play a major role in the stabilization of biofilm architecture by connecting bacterial cells to the EPS matrix [Fong and Yildiz, 2015]. Among the structural proteins, lectins represent an important subclass that mediate cell–matrix and cell–cell interactions through specific carbohydrate recognition and therefore play an important role in establishing biofilms [Napoleão et al., 2025].

*P. aeruginosa* is a major threat to clinical settings and plays a detrimental role in a large number of hospital-acquired infections (HAIs) [Moradali et al., 2017; Reynolds and Kollef, 2021]. It tends to cause multi-site infections, of which bacteremia is fatal, with a mortality rate ranging from 18% - 61% [Zhang et al., 2020]. LecA and LecB are the carbohydrate-binding proteins (lectins) present in the EPS matrix of *P. aeruginosa* biofilms that play a major role in maintaining its architecture. LecA shows high binding affinity with galactose and its derivatives [Garber et al. 1992]. In acute infections, LecA mediates the initial interaction with the host cells by acting as an adhesion factor, whereas in the case of chronic infections, it contributes to biofilm formation by *P. aeruginosa* [Chemani et al. 2009]. On the other hand, LecB contributes to biofilm formation by binding to Psl, a key polysaccharide present in the EPS matrix of *P. aeruginosa* biofilms. The above interaction between the LecB protein and Psl polysaccharide leads to increased retention of both bacterial cells and EPS in a developing biofilm. Therefore, it is evident that biofilm formation and associated virulence in *P. aeruginosa* can be prevented by inhibiting the function of the above lectins [Wagner et al., 2017].

Specific targeting by small molecules could be a potent strategy in order to disintegrate bacterial biofilms. In this context, researchers have often exploited nanoscale molecules such as nanoparticles (NPs), nanoclusters, and carbon dots (Cdots) [Amod et al., 2025; Verma et al., 2020]. Apart from nanomaterials, antimicrobial peptides (AMPs) are another class of small molecules that can be leveraged to target bacterial biofilms. These are small peptide molecules, less than 50 amino acids in length, that are endogenously produced and released in almost all living organisms as a part of their innate immune response [Joo et al., 2016]. AMPs have been reported to show their antibacterial activity through several mechanisms, which include disruption of the cell membrane, binding to nucleic acid, among others [Singh et al., 2023]. Therefore, unlike antibiotics, the AMPs have a different set of mechanisms to target the same bacteria, which reduces the chances of bacteria to evade their bactericidal action [Yasir et al., 2018]. Importantly, AMPs show high collateral sensitivity and low cross-resistance, which makes them more promising as antimicrobial agents [Lázár et al. 2019]. They can be used in combination with other antibacterial and antibiofilm agents to achieve enhanced efficacy against bacteria and their biofilms [Anand et al., 2024a]. Additionally, AMPs are reported to possess enormous potential to exhibit prominent antibacterial and antibiofilm activity against different drug-resistant bacteria. Researchers have also started exploring ultrashort peptides (USPs) as antibacterial and antibiofilm agents. They are functional derivatives of full-length AMP, 3-8 amino acids long [Almaaytah et al., 2018]. USPs have the added advantage of lower cost of synthesis with increased translationary potential [Afami et al., 2021]. In addition, USPs are also expected to show better biocomptability in relation to the parent AMPs

The human gut microbiome is a rich reservoir of AMPs and is involved in several activities such as immunomodulation, disruption of bacterial cell membrane and halting nucleic acid synthesis [Cotter et al., 2013]. As compared to other sources, AMPs tapped from human gut microbiota are expected to show low toxicity and immunogenicity since the commensal gut bacteria producing the AMPs are adapted to coexist within the human gastrointestinal environment over the course of millions of years of evolution [Dominguez-Bello et al., 2019].

Keeping the above fact in mind, in our current study, we first developed a comprehensive *in-silico* pipeline to identify putative antibacterial and antibiofilm AMPs present in the human gut microbiota. Subsequently, we performed molecular docking and molecular dynamics (MD) simulation studies to determine the interaction of these AMPs with the lectin proteins of *P. aeruginosa* (i.e., LecA and LecB). Among the selected peptides, amp6, amp21, and amp24 were found to be the most potent inhibitors of the above lectin proteins and were therefore selected for *in-vitro* validation. The amp21 showed excellent biocompatibility and the most prominent antibacterial activity. Biofilm inhibition and disruption assays showed that amp6 and amp21 brought a greater reduction in the biofilm mass of *P. aeruginosa* when compared to amp24. All three selected AMPs showed significant antibiofilm activity by stimulating the transition of bacterial cells from the biofilm to the free-living planktonic state. It is conceivable that the above-observed phenomenon was due to the interaction of the AMPs with the lectin proteins, which subsequently disturbed the integrity of the EPS matrix. In addition, the AMPs exhibited antibacterial activity by causing perturbations in the bacterial cell membrane.

## 2. Materials and methods

### 2.1. Materials

Luria Bertani (LB) broth, Muller Hinton agar (MHA), Dimethyl sulfoxide (DMSO), 99% glacial acetic acid, sodium chloride, magnesium sulphate, L-arginine, potassium phosphate, streptomycin sulphate, and tetracycline hydrochloride were procured from SRL, India. Propidium iodide (PI), trypsin, and 3-(4,5-dimethylthiazol-2-yl)-2,5-diphenyltetrazolium bromide (MTT) and RPMI 1640 media were purchased from Sigma-Aldrich. Phosphate buffer saline (PBS) and Triton X-100 were bought from Himedia, India. Antibiotic-antimycotic mix and Foetal bovine serum (FBS) were purchased from GIBCO. Glucose was purchased from Merck. *Pseudomonas aeruginosa* PAO1 strain was a kind gift from Prof. SomLata from Jamia Milia Hamdard, New Delhi, India. The HCT116 cancer line was procured from the National Centre for Cell Science (NCCS), Pune. The lead AMPs filtered from the *in-silico* studies were sent for synthesis to the Peptide Purchase Committee, Elabscience, USA. The AMPs were supplied as lyophilized powder and were stored in -20 °C post receipt.

### 2.2. Methods

#### 2.2.1. Evolutionary study of lectin proteins

Before targeting the LecA and LecB proteins of *P. aeruginosa*, their evolutionary lineage was studied through phylogenetic analysis with the objective of (i) estimating the degree of structural conservation of these lectins among other ESKAPE pathogens (ii) determining the presence of their structural homologs in humans, to gain insights into possible cross-reactivity or toxic effects while targetting these lectins.

For this purpose, BlastP (https://blast.ncbi.nlm.nih.gov/Blast.cgi?PAGE=Proteins) analysis was performed on NCBI using the FASTA sequences of LecA (PDB ID: 6YOH) and LecB (PDB ID: 1OUX) proteins [Camacho et al., 2009]. The number of hits found for LecA was 742, whereas for LecB was 1520. The FASTA sequences of the BLASTP hits were downloaded and imported into the MEGA v11 software [Tamura et al., 2021]. Sequences belonging to the ESKAPE group were selected for further analysis using the following keywords: *‘Escherichia coli,’ ‘Staphylococcus aureus,’ ‘Klebsiella pneumonia,’ ‘Acinetobacter baumannii,’ ‘*Pseudomonas,’ ‘*Enterococcus faecium,’ ‘*Enterobacter,’ and ‘Enterobacteriaceae.’

Multiple sequence alignment (MSA) was performed using MUSCLE (https://www.ebi.ac.uk/jdispatcher/msa/muscle?stype=protein) [Edgar, 2004]. The phylogenetic tree was constructed using the Maximum-Likelihood method, and the tree was exported in Newick format for visualization. Finally, the phylogenetic tree was visualized and annotated using iTOL software (https://itol.embl.de/) [Letunic and Bork, 2021].

#### 2.2.2. Selection of AMPs for screening

After the evolutionary conservation and host-specificity insights were obtained for LecA and LecB proteins through phylogenetic analysis, the next step involved the rational selection of AMPs for screening to target the above lectin proteins. The AMPs used in this study were taken from the study by Anand et al. (2025), where a comprehensive *in-silico* pipeline was developed to identify and characterize putative antibacterial and antibiofilm peptides produced by human gut microbiota [Anand et al., 2025].

#### 2.2.3. Retrieval of target-protein structures

Once the AMPs were selected, the 3-D X-ray crystallographic structures of the target proteins, i.e., LecA and LecB (from *P. aeruginosa*), were downloaded from the PDB database (https://www.rcsb.org/) [Berman, 2000].

Structures of LecA protein with the PDB IDs: 7Z62/7Z63, 5MIH, and 4LK6/4LK7/4LKE either represented different chemotypes or lacked the specific inhibitor context required for docking optimization [Bruneau et al., 2023; Wagner et al., 2017; Kadam et al., 2013]. LecA structure with the PDB ID (6YOH) was selected for the docking studies owing to its high-resolution structure (1.84 Å) and ligand-bound conformation with a relevant glycomimetic inhibitor [Kuhaudomlarp et al., 2020].

On the other hand, structures of LecB protein with the PDB IDs: 5A6X, 5A6Y, 5A6Z, 5A70, and 5MB1reflected different ligand chemotypes and subtle sequence divergences and were therefore not suitable to use in a uniform docking framework [Lepsik et al., 2019; Sommer et al., 2016; Sommer et al., 2018]. LecB protein with the PDB ID 1OUX was selected for the docking studies since it provided a standard reference scaffold with well-characterized fucose/mannose binding fold [Loris et al., 2003].

#### 2.2.4. Three-dimensional modelling of AMPs

The 3-D modelling of AMPs was performed using the PEP-FOLD3.5 software as per the protocol mentioned by Anand et al. (2025) [Anand et al., 2025].

#### 2.2.5. Molecular docking and MM/GBSA analysis

Molecular docking and MM/GBSA analyses were performed between the selected AMPs and the 3D structures of the lectin proteins on HawkDock server (http://cadd.zju.edu.cn/hawkdock/) to evaluate their binding interactions [Weng et al., 2019]. The MM-GBSA binding free energy gives a more accurate assessment of the thermodynamic stability of each complex by considering solvation and entropic effects, whilst the docking score gives an indication of the overall binding affinity and interaction strength between the target protein and the peptides. Lower (more negative values) for both parameters indicates more stable and stronger binding between the peptides and the target protein. Further, in order to determine the contribution of each residue to the total binding free energy, MM/GBSA free energy decomposition analysis and residue analysis were performed [Dalal et al., 2021; Kumari and Dalal, 2022].

#### 2.2.6. MD simulation analysis

MD Simulation was carried out for a duration of 100 ns using the protocol mentioned by Anand et al., 2024 [Anand et al., 2024b].

#### 2.2.7. Analysis of MD Simulation results

GROMACS utilities were employed to obtain the trajectories for performing the MD simulation studies. For this objective, Root mean square deviation (RMSD), radius of gyration (Rg), hydrogen bond interactions, coul-SR and LJ-SR interactions were measured. Free energy landscape (FEL) plots were generated by utilising gmx-sham [Prakash et al., 2018].

#### 2.2.8. Biocompatibility of the lead AMPs

Before studying the antibacterial and antibiofilm activity of the lead AMPs (screened through MD simulation studies), they were tested for their biocompatibility through the hemocompatibility and cytotoxicity assay:

##### 2.2.8.1. Hemocompatibility assay

The hemocompatibility of the AMPs was investigated in accordance with our previously standardized protocol [Singh et al., 2022]. The human red blood cells (RBCs) were mixed with AMPs at varying concentrations between 6.25-200 µg mL^-1^. Hemolytic activity of AMPs was expressed in percentage relative to the hemolysis induced by Triton X-100, which was considered 100% that was used as a positive control. PBS served as the negative control. The experiment was performed twice independently (in triplicate), and the results are represented as the mean of both times.

##### 2.2.8.2. Cytotoxicity assay

Post hemocompatibility assessment, the cytotoxic effects of the lead AMPs were further evaluated in mammalian cell lines through MTT dye reduction assay to evaluate their cellular safety profile. In brief, a confluent monolayer of human colorectal cancer (HCT116) cells was trypsinized using (0.25% Trypsin - 0.1% EDTA), following which the cells were harvested by centrifugation. The harvested cells were then seeded in a 96-well plate at a density of 3 × 10^3^ cells per well in 100 μL of complete RPMI1640 media (RPMI1640 supplemented with FBS and antibiotic-antimycotic mix). The cells were allowed to grow at 37 °C for 24 h to permit cell attachment and further growth. The medium was then removed and replaced with 100 µL of fresh medium with or without AMPs (at varying concentrations ranging between 6.25 µg mL^-1^ to 100 µg mL^-1^). After treatment, 10 µL of 5 mg ml^−1^ MTT solution was added to each well and was further incubated for an additional 4 h at 37 °C. The supernatant was carefully removed, and the formazan crystals formed were dissolved in 100 µL of DMSO. The OD was estimated through a microplate reader at 570 nm, and the viability of HCT116 cells against the peptides was determined in terms of percentage. The viability of the HCT116cells treated with AMPs was expressed in percentage relative to the viability of the untreated cells, which was considered 100%. DMSO was used as the blank. The experiment was performed twice independently (in triplicate), and the results are represented as the mean of both times.

#### 2.2.9. Antibacterial efficacy of AMPs against *P. aeruginosa*

Once the biocompatibility of the lead AMPs was determined, we subsequently investigated their antibacterial potency. Firstly, in accordance with the CLSI guidelines, we first evaluated the MIC of the lead AMPs against *P. aeruginosa* using the standard broth microdilution method, ranging from 0.25 to 64 µg mL^-1^ for 12 h. However, we did not obtain MIC up to 64 µg mL□¹.

Therefore, subsequently, we performed a CFU-based viability assay to evaluate concentration-dependent reductions in bacterial load upon treatment with the lead AMPs. Stationary phase cells of *P. aeruginosa* were treated with the AMPs at concentrations of 50 µg mL^-1^ and 100 µg mL^-1^. These were then incubated at 37 °C for a duration of 24 h under static conditions. Before treatment, *P.aeruginosa* culture was standardized to an initial OD□□□ of 0.002. Subsequently, the untreated and *P. aeruginosa* cells treated with AMPs were diluted 10^-6^ times. 100 µL of dilution of all the samples was then spread uniformly on MHA plates and incubated at 37 °C for 24 h. The CFU mL^-1^ was calculated by the following equation:

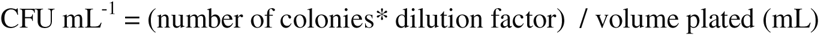

In this experiment, the untreated *P. aeruginosa* cells were considered to be the control, and the bacterial load in the treated sample was plotted as % CFU mL^-1^ with respect to the untreated control. The CFU mL^-1^ in the untreated *P. aeruginosa* cells were considered to be 100%. The experiment was performed twice, and the results are represented as the mean of both times.

#### 2.2.10. PI uptake assay

The fluorescence of propidium iodide (PI) was measured in *P. aeruginosa* treated with lead AMPs to determine whether membrane permeabilization contributed to the antibacterial activity of the AMPs. An increase in PI fluorescence corresponds to increased cell membrane permeability. For this purpose, an overnight culture of *P. aeruginosa* was incubated with lead AMPs at concentrations of 12.5 µg mL^-1^ and 100 µg^-1^ mL for 3 h at 37 °C under shaking conditions. Following incubation, the samples were centrifuged at 8000 rpm for 5 min, washed with 1X PBS, and centrifuged again under identical conditions. The supernatants were discarded, and the cell pellets were resuspended in 1X PBS containing 10 µg mL^-1^ PI. Fluorescence measurements for all the samples were recorded at an excitation wavelength of 488 nm. Herein, the untreated (control) *P. aeruginosa* cells served as the negative control.

#### 2.2.11. Antibiofilm efficacy of AMPs against *P. aeruginosa* biofilms

Following the evaluation of antibacterial activity of the lead AMPs, their antibiofilm potential was examined against *P. aeruginosa* through both biofilm inhibition and biofilm disruption studies:

##### 2.2.11.1 Inhibition of *P. aeruginosa* biofilm by AMPs

For studying the biofilm inhibition efficacy of the lead AMPs, stationary phase cells of *P. aeruginosa* (grown in LB broth) were treated with 50 µg mL^-1^ of all the lead AMPs. These were then incubated in polystyrene-coated 96-well plates under static conditions at 37 °C for 24 h. Herein, the *P. aeruginosa* inoculum was standardized to an initial OD□□□ of 0.02 while setting up the biofilm. The untreated wells were taken as the control. Post treatment, the media was carefully aspirated without disturbing the biofilm and allowed to air-dry for 30 min. To fix the biofilm, 200 µL of methanol was added to each well and incubated at room temperature for 30 min. Following this, the media was removed, and the plate was kept for drying for 2 h. Subsequently, 0.1% (w/v) crystal violet solution was used to stain the biofilms. The staining solution was removed after 10 min of incubation at room temperature. In order to solubilize the bound dye, 30% glacial acetic acid was added, and the 96-well plate was again incubated for 10 min at room temperature. Relative quantification of biofilm mass was performed through a microplate reader by measuring the optical density at 540 nm. The mass of the AMP-treated biofilms was expressed in percentage relative to the biomass of untreated biofilms, which was considered to be 100%. The experiments were performed thrice independently (in triplicate), and the data were plotted as mean ± SD.

##### 2.2.11.2 Disruption of *P. aeruginosa* biofilm by AMPs

For studying the biofilm disruption efficacy of the lead AMPs, mature biofilms were first obtained by incubating stationary phase cells of *P. aeruginosa* (grown in LB broth) in polystyrene-coated 96-well plates under static conditions at 37 °C for 48 h. Herein, the *P. aeruginosa* inoculum was standardized to an initial OD□□□ of 0.02 while setting up the biofilm. The above biofilm was then treated with 50 µg mL^−1^ of each of the lead AMPs and was further incubated for 24 h under the same conditions. Post treatment, the biofilm mass was quantitatively estimated in accordance with the standard protocol mentioned in section 2.2.11.1. The experiments were performed thrice independently (in triplicate), and the data were plotted as mean ± SD.

#### 2.2.12. SEM analysis to study morphology of *P. aeruginosa* biofilm

Scanning electron microscopy (SEM) was used to observe the effects of lead AMPs on biofilm integrity and other morphological changes in the bacterial cells. For biofilm formation, sterile circular coverslips (18 mm diameter) were placed in the wells of a flat-bottom 6-well polystyrene tissue culture plate, and *P. aeruginosa* biofilms were allowed to develop over them.

Herein, the *P. aeruginosa* inoculum was standardized to an initial OD□□□ of 0.02 while setting up the biofilm. Herein, the 48 h biofilm of *P. aeruginosa* was treated with 100 µg mL^-1^ of amp6 (as a representative) and was further incubated at 37 °C for 12 h. Post treatment, biofilms were fixed with 4% PFA for 5-10 min at room temperature and were then kept overnight at 4 °C. On the following day, biofilm samples were dehydrated with a graded series (30%, 50%, 70%, 90%, and 100%) of ethanol for 5 min each on a gel rocker. Eventually, the biofilms were coated by a gold sputter coater and examined using a FEI Quanta 250 Scanning Electron Microscope.

#### 2.2.13. Mechanistic evaluation of antibiofilm activity

In order to decipher the mode of mechanism for the antibiofilm activity of the AMPs, mature biofilms were first obtained by growing stationary phase cells of *P. aeruginosa* at 37 °C for 48 h in test-tubes under static conditions. Herein, the *P. aeruginosa* inoculum was standardized to an initial OD□□□ of 0.02 while setting up the biofilm. The above biofilm was then treated with 200 µg mL^−1^ of all the lead AMPs and was further incubated for 24 h under the same conditions. Subsequently, the integrity of the biofilm pellicle was visually observed to study the biofilm disruption by AMPs. Post treatment, a CFU assay was performed for all the samples using the planktonic cells present in the media beneath the biofilm pellicle. The planktonic suspensions aliquoted from the media (beneath the biofilm pellicle) were diluted 10^-6^ times. These were then spread uniformly on MHA plates and incubated at 37 °C for 24 h. Herein, the untreated *P. aeruginosa* biofilm was considered to be the control. The experiments were performed thrice independently in triplicate.

#### 2.2.14. Derivation of USPs from the lead AMPs and their interaction with LecA protein

USPs were derived from the lead AMPs targeting the LecA protein. Initially, key residues of the lead AMPs targeting the LecA protein were identified through residue interaction analysis using the previously obtained MD simulation data. After identification of the key binding residues, USPs were manually designed from the lead AMPs. The interaction of these USPs with the LecA protein was then studied through molecular docking, MMGBSA and MD simulation analysis.

##### 2.2.14.1. Residue interaction analysis

For residue interaction analysis, the centered trajectory files from the 100 ns MD simulations performed between LecA protein and the lead AMPs were utilized. A complex PDB file with the ensemble of structures created during the simulation period was taken from these trajectories. In order to find and examine non-covalent interactions within the complexes, this PDB file was subsequently uploaded to the Residue Interaction Network Generator (RING) server (https://ring.biocomputingup.it/) [Del Conte et al., 2024]. The RING software uses particular geometric thresholds to identify different kinds of residue interactions.

Based on certain geometric thresholds, the RING software finds different kinds of residue interactions. For this study, the following distance criteria were used: 3.9 Å for the hydrogen bond donor–acceptor distance, 2.5 Å for the hydrogen bond H–acceptor distance, 4.0 Å for the ionic bond distance, 5.0 Å for the π–cation distance, 6.5 Å for the π–π stacking center distance, 4.3 Å for the π–hydrogen donor ring center distance, 2.8 Å for the metal ion coordination distance, and 2.5 Å for the disulfide bond distance. The RING output gives interaction probability scores between 0 and 1, which show how likely the interaction will happen during the simulation. After that, these scores were turned into frequency percentages to make it easier to compare them. Finally, the interactions between the protein and peptide chains were plotted to show which residue of the protein interacts with which residue of the peptide.

##### 2.2.14.2. Designing of USPs

To design the USPs, the interaction profiles between the lead peptides and the LecA protein were analyzed in detail. The amino acid residues of the peptide that directly interact with particular residues of LecA were identified using the residue-level interaction data derived from the RING analysis. In particular, peptide residues that formed interactions with the galactose-, glucose-, or fructose-binding moieties of the LecA protein were identified and prioritized for further design.

Based on these interaction sites, new USP sequences were manually designed. The approach involved retaining only those regions of the parent peptides that contained residues showing direct interaction with the carbohydrate-binding residues of LecA. Each selected fragment was then truncated or extended slightly to ensure structural continuity and preserve key interaction motifs.

The final designed peptides varied in length from 5-8 amino acids, representing the minimal motifs essential for stable interaction with LecA.

##### 2.2.14.3. Peptide folding and structure modeling of USPs

All of the manually designed USPs were modelled using the PEP-FOLD3 server (http://bioserv.rpbs.univ-paris-diderot.fr/services/PEP-FOLD3). This server performs *de novo* free or biased prediction for linear peptides 5-50 amino acids long. Further, it can also generate native-like conformations of a peptide interacting with a protein provided the interaction site is known in advance. [Lamiable et al., 2016].

##### 2.2.14.4. Molecular docking and MM/GBSA analysis

Molecular docking and binding free energy estimation were carried out using the HawkDock server (http://cadd.zju.edu.cn/hawkdock/). The analysis was performed between the LecA protein and the designed USPs. The docking results were subsequently refined and evaluated using MM/GBSA calculations to estimate the binding free energy of the protein–peptide complexes. For the 2D interaction analysis of the docked complexes, we used the Ligplot+ v2.3 software [Laskowski and Swindells, 2011].

##### 2.2.14.5. MD simulation

The top-scoring LecA–USP complexes were selected for MD simulations in GROMACS v2019.4 following the completion of the docking studies. To see how the complex behaved over time and whether the interactions were stable, each system was operated for 200 ns in an explicit solvent environment. The force field and other simulation parameters were set up using the same process as performed by Anand et al. 2024b [Anand et al., 2024b].

##### 2.2.14.6. Protein–Peptide MD simulation analysis

To analyze the generated trajectories, post-simulation evaluations were conducted utilizing various GROMACS tools. We used the RMSD, Rg, solvent-accessible surface area (SASA) and root-mean-square fluctuation (RMSF) to figure out how stable and flexible the USP-LecA complexes were. To see the conformational energy states of the systems, the gmx sham module was used to make the FEL plots [Prakash et al., 2018].

## 3. Results and discussion

### 3.1. Homology search and evolutionary study of lectin proteins

The complete BLASTP output for LecA and LecB can be studied from the supplementary information (Supplementary file 1). A comparative sequence and phylogenetic analysis of LecA and LecB was performed to evaluate the evolutionary conservation and target specificity of lectin proteins across clinically relevant pathogens. The initial BlastP analysis confirmed that both LecA and LecB proteins of *P. aeruginosa* have no structural homologs in humans. This implies that targeting these lectin proteins with any antimicrobial molecule will only affect *P. aeruginosa*, thereby minimizing the risk of off-target toxicity in humans and improving their safety for therapeutic uses.

Representative or dummy trees were built such that the positioning of *P. aeruginosa* with respect to other bacteria remains least affected. The same can be visualized in Figure 1a-b. The phylogenetic tree of LecA revealed that *P. aeruginosa* clusters closely with *K. pneumoniae*, suggesting that these two ESKAPE pathogens share a high level of sequence conservation for the LecA protein, as visible in the representative tree in Figure 1a. Given their close evolutionary relationship, it is possible that the lectin domains involved in adhesion and carbohydrate recognition may remain structurally intact and continue to play comparable functions in host-pathogen interactions. On the other hand, different clades were formed by Enterobacter species (such as *E. cloacae*, *E. hormachei*, and *E. ludwigii*), which suggested that the LecA orthologs of these strains differ from one another. Certain Enterobacter subgroups exhibited modest branch length values (7.1e–7), which reflected minor sequence changes rather than total functional loss. According to this conservation pattern, AMPs made to target LecA may be effective against several pathogens, including *P. aeruginosa* and *K. pneumoniae*, but they may also have varying binding efficiencies in Enterobacter species because of structural or sequence differences.

**Figure 1.**
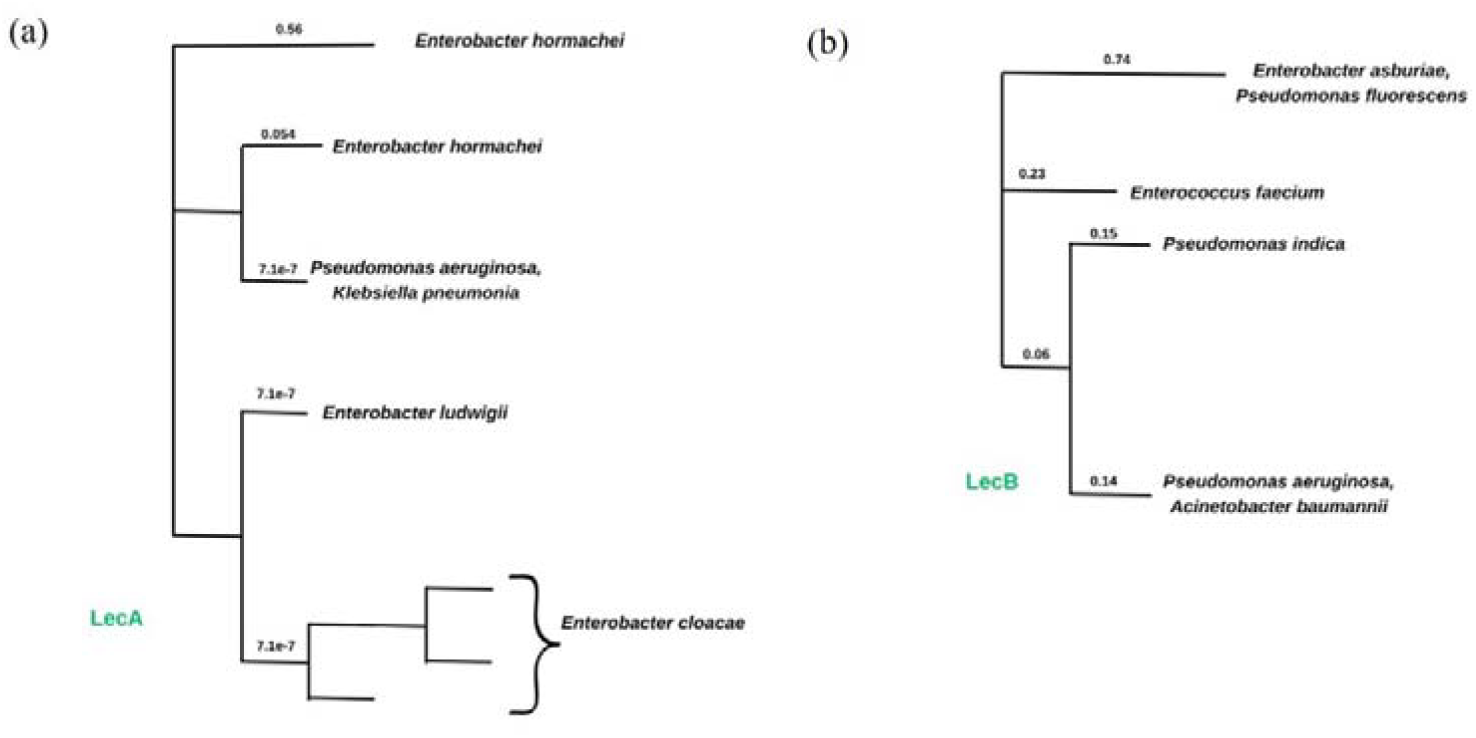
(a) Representative image for phylogenetic tree of LecA protein among ESKAPE pathogens **(b)** Representative image for phylogenetic tree of LecA protein among ESKAPE pathogens

The phylogenetic tree of LecB showed that it exhibits a high degree of sequence conservation in *P. aeruginosa* and *A. baumannii* (branch length 0.14), which reflected their evolutionary proximity (Figure 1b). Therefore, the AMPs targeting the LecB protein in *P. aeruginosa* are expected to exhibit cross-reactivity against *A. baumannii*, another clinically significant ESKAPE pathogen. On the other hand, *Pseudomonas indica* and *Enterococcus faecium* formed distinct, distantly related branches, which suggested significant sequence divergence and potentially modified carbohydrate-binding motifs. *Pseudomonas fluorescens* and *Enterobacter asburiae* were part of the most diverse cluster; their comparatively long branches (0.74) indicated less evolutionary conservation. The effectiveness of AMP in targeting LecB was found to be pathogen-dependent. It may be strong against *Pseudomonas* and *Acinetobacter*.

### 3.2. Selection of gut-derived AMPs

Prospective antibacterial and antibiofilm gut-derived peptides identified by Anand et al. (2025) were selected for the study [Anand et al., 2025]. The sequences of the 24 gut-derived AMPs were obtained from the same research group. Consequently, the sequences of the AMPs have been displayed in Table 1.

**Table 1.**
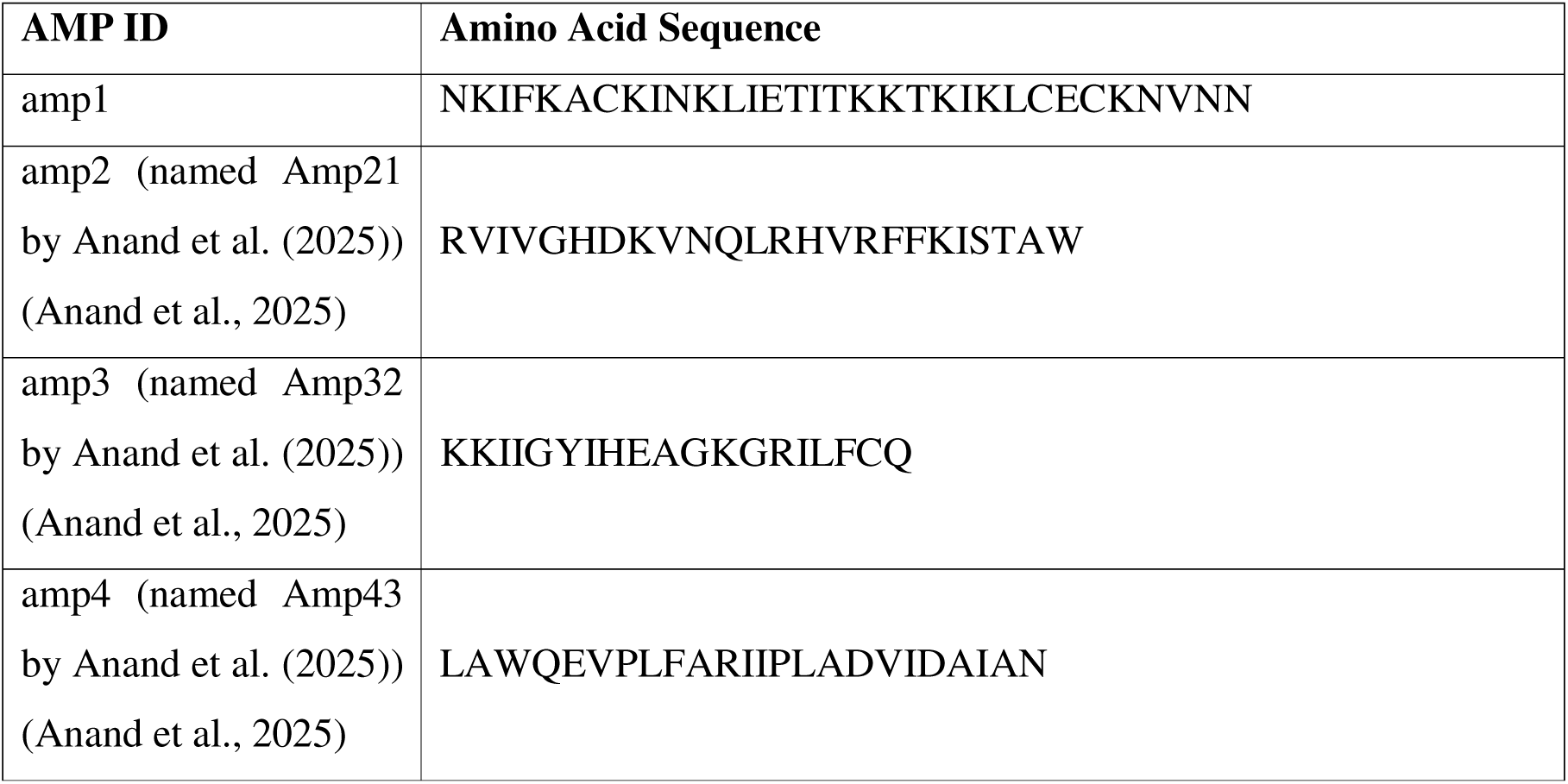
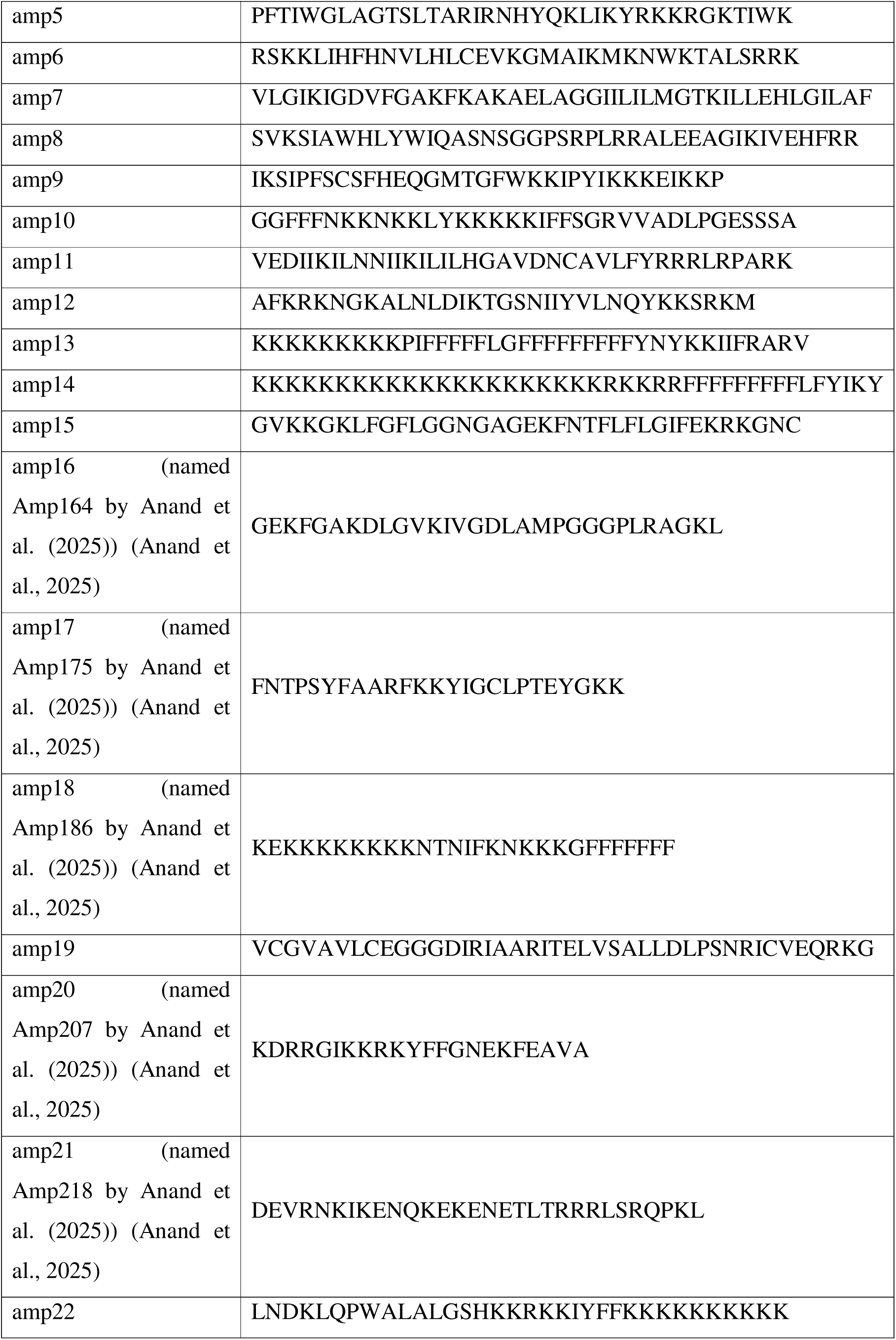
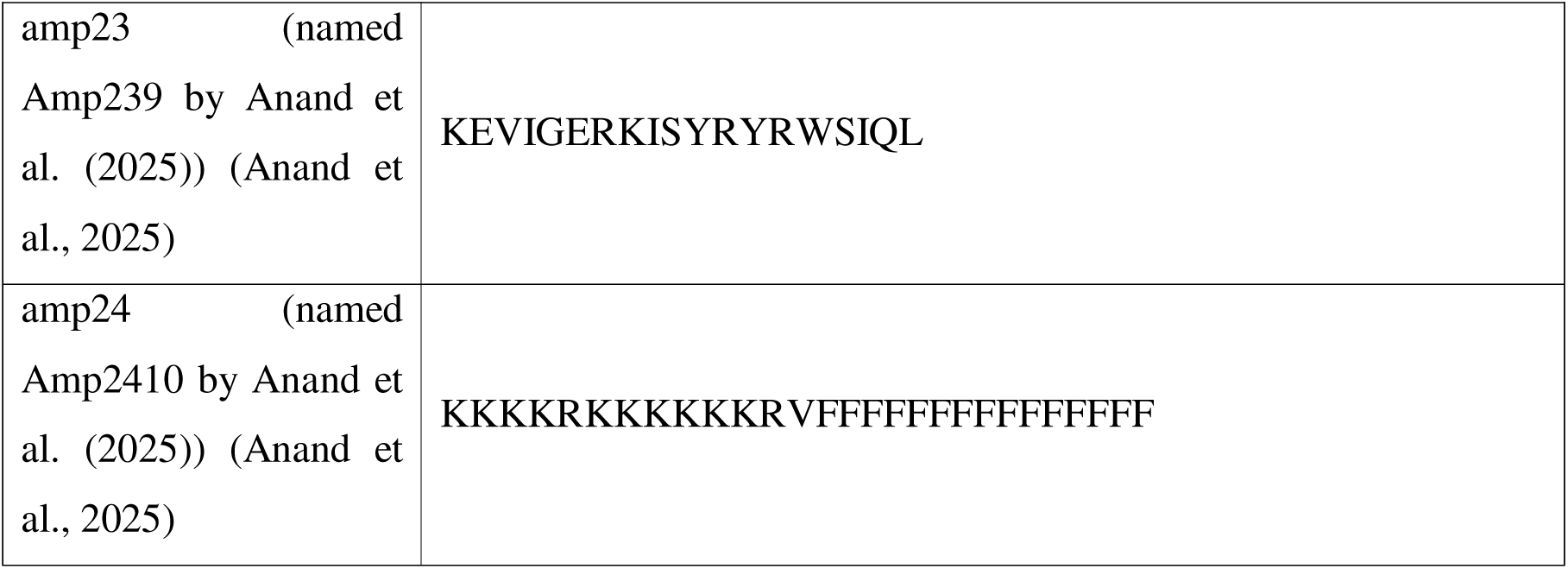
Sequence of gut-derived AMPs.

### 3.3. 3D modelling of the peptides on PEP-FOLD3

Following the phylogenetic analysis of the lectin proteins, the 3D structures of the AMPs identified through the *in silico* pipeline were modelled using PEP-FOLD3 and visualized in PyMOL. All the peptides showed a predominant α-helical conformation.

### 3.4. Studying the binding affinity of the lead AMPs with the lectin proteins through molecular docking and MM/GBSA analysis

Among all the AMPs, amp14 showed the best binding profile with LecA protein with a docking score of -4374.23 and an MM-GBSA of -85.03 kcal mol□¹. This indicated the formation of a very stable complex with LecA. Following this, energetically favourable interactions were obtained for amp13–LecA (MMGBSA:–60.78 kcal mol□¹; docking score: –3743.71), amp22–LecA (MMGBSA: –60.18 kcal mol□¹; docking score: –3556.73), and amp24–LecA (MMGBSA: –52.20 kcal mol□¹; docking score: –2974.60) complexes. Interestingly, amp8–LecA complex showed a highly negative docking score (–4063.08), indicating significant form complementarity and robust binding at the LecA interface, despite having a relatively higher MM-GBSA (–43.11 kcal mol□¹). The binding energy and docking score were also found to be favourably balanced in amp6–LecA (MMGBSA: –49.32 kcal mol□¹; docking score:–3982.64) and amp21–LecA (MMGBSA: –48.97 kcal mol□¹; docking score: –3820.95) complexes. To further evaluate their structural stability and dynamic behavior under physiological conditions, the seven complexes: amp14–LecA, amp13–LecA, amp22–LecA, amp24–LecA, amp6–LecA, amp21–LecA, and amp8–LecA were selected for molecular dynamics (MD) simulation studies based on thermodynamic stability and interfacial interaction strength. The MM/GBSA and Hawdock docking scores of LecA-AMP complexes have been listed in Table 2.

**Table 2.** MM/GBSA and Hawkdock docking score of LecA-AMP complexes.

| **Complex** | **MMGBSA- Binding Free Energy (kcal mol<sup>-1</sup>)** | **Hawkdock docking score** |
| --- | --- | --- |
| amp14-LecA | -85.03 | -4374.23 |
| amp13-LecA | -60.78 | -3743.71 |
| amp22-LecA | -60.18 | -3556.73 |
| amp24-LecA | -52.2 | -2974.6 |
| amp20-LecA | -49.66 | -2796.48 |
| amp18-LecA | -49.61 | -3402.78 |
| amp6-LecA | -49.32 | -3982.64 |
| amp21-LecA | -48.97 | -3820.95 |
| amp15-LecA | -48.27 | -3094.87 |
| amp11-LecA | -47.87 | -3230.46 |
| amp5-LecA | -47.74 | -2872.6 |
| amp17-LecA | -46.06 | -3336.94 |
| amp7-LecA | -44.99 | -3436.61 |
| amp19-LecA | -44.68 | -3027.59 |
| amp2-LecA | -43.73 | -3638.95 |
| amp8-LecA | -43.11 | -4063.08 |
| amp9-LecA | -43.11 | -3149.36 |
| amp10-LecA | -40.98 | -3327.84 |
| amp12-LecA | -40.87 | -3531.43 |
| amp23-LecA | -37.65 | -2601.07 |
| amp16-LecA | -37.28 | -2491.12 |
| amp3-LecA | -35.37 | -2842.58 |
| amp4-LecA | -33.82 | -3373.51 |
| amp1-LecA | -33.02 | -3389.97 |

With a docking score of -4554.76 and a MM-GBSA binding free energy of -115.42 kcal mol□¹, amp14 demonstrated excellent thermodynamic stability on interaction with the LecB protein. The docking scores of amp22–LecB (MMGBSA: –98.64 kcal mol□¹; docking score: –5192.46), amp13–LecB (MMGBSA: –92.03 kcal mol□¹; –5251.25), and amp11–LecB (–86.27 kcal mol□¹; –4863.99) complexes were similarly strong or even better, indicating improved interfacial complementarity and the possibility of successful binding. Furthermore, although having somewhat higher MM-GBSA energies, amp6–LecB (MMGBSA: –73.29 kcal mol□¹; docking score: –5881.09) and amp21–LecB (MMGBSA: –59.92 kcal mol□¹; docking score: –5691.76) complexes showed unusually negative docking scores, which indicated strong peptide-lectin interactions. Finally, six complexes, i.e., amp14–LecB, amp22–LecB, amp13–LecB, amp11–LecB, amp6–LecB, and amp21–LecB, were chosen for MD simulations in order to assess their conformational stability, flexibility, and interaction persistence under physiological conditions. This was done after taking into account both energetic stability and binding affinity. The MM/GBSA and Hawdock docking scores of LecB-AMP complexes have been listed in Table 3.

**Table 3.** MM/GBSA and Hawkdock docking score of LecB-AMP complexes.

| **Complex** | **MMGBSA- Binding Free Energy**<br>**(kcal mol<sup>-1</sup>)** | **Hawkdock docking score** |
| --- | --- | --- |
| amp14-LecB | -115.42 | -4554.76 |
| amp22-LecB | -98.64 | -5192.46 |
| amp13-LecB | -92.03 | -5251.25 |
| amp11-LecB | -86.27 | -4863.99 |
| amp24-LecB | -84.72 | -4896.87 |
| amp18-LecB | -84.3 | -4547.95 |
| amp10-LecB | -76.25 | -4305.42 |
| amp6-LecB | -73.29 | -5881.09 |
| amp2-LecB | -70.6 | -4297.64 |
| amp12-LecB | -64.4 | -4305.42 |
| amp3-LecB | -63.16 | -3666.27 |
| amp9-LecB | -62.71 | -3885.85 |
| amp8-LecB | -60.74 | -4301.94 |
| amp21-LecB | -59.92 | -5691.76 |
| amp20-LecB | -57.28 | -3832.56 |
| amp1-LecB | -56.19 | -3875.4 |
| amp7-LecB | -56.17 | -4301.94 |
| amp16-LecB | -55.85 | -3644.46 |
| amp5-LecB | -55.14 | -3592.51 |
| amp15-LecB | -50.41 | -3667.41 |
| amp23-LecB | -50.37 | -3393.23 |
| amp4-LecB | -47.55 | -3315.72 |
| amp17-LecB | -46.09 | -3500.11 |
| amp19-LecB | -36.89 | -3992.06 |

### 3.5. Assessment of the conformational stability and binding behavior of top-ranked AMP–lectin complexes through MD simulation analysis

To assess their conformational stability and binding behavior of top-ranked AMP–lectin complexes obtained from molecular docking, MD simulations were carried out. The simulations were evaluated using RMSD, Rg, Coul-SR and LJ-SR interaction energies, H-bonding patterns, and FEL analysis, which provide key insights into the stability and energetically favorable conformations of the complexes.

#### 3.5.1. Analyzing the dynamic stability and interaction profiling of LecA–peptide complexes

RMSD measures the average deviation in atomic position of a structure (often the backbone or all heavy atoms) from a reference structure over time during MD simulation. Out of all the LecA-peptide complexes, amp21-LecA showed the lowest RMSD. This led to an inference that, with respect to the other AMPs, amp21 remains bounded in a constant orientation at the binding pocket of the LecA protein (Figure 2a). Following this, Rg was measured in order to assess the compactness and folding behaviour of LecA-peptide complexes over the duration of simulation. Herein, amp24-LecA showed a low and constant Rg among all the LecA-peptide complexes. This suggested that (i) binding of the amp24 with LecA led to structural tightening and compaction, (ii) the amp24-LecA complex maintained a comparatively more consistent and compact structure during MD simulation with respect to other complexes (Figure 2b). The short-range electrostatic interaction energy between the protein and peptide is represented by Coul-SR, which measures how strongly their charged or polar residues interact at a typical cutoff distance of ≤1.0–1.2 nm. Among all the LecA-peptide complexes, amp14-LecA showed the most-negative Coul-SR, which implied that the electrostatic interactions between amp14 and the LecA protein are the strongest when compared to other AMPs (Figure 2c). The short-range van der Waals or non-polar interaction energy between the protein and peptide atoms is indicated by LJ-SR, which shows how closely packed residues interact within a typical cutoff distance of ≤1.0–1.2 nm. In our study, it was observed that the amp13-LecA complex showed the lowest LJ-SR values among other LecA-peptide complexes. This observation led to an inference that among all the AMPs, amp13 showed more pronounced hydrophobic /Van der Waals interactions with the LecA protein (Figure 2d). A higher number of H-bonds in protein-peptide simulation indicates greater stability of the protein-peptide complex. The stability, strength, and specificity of the protein–peptide interaction can be assessed by counting the hydrogen bonds during MD simulation, which also provides information on the complex’s structural integrity and binding affinity. It was found that amp14 and amp22 formed a greater number of H-bonds with the LecA protein. However, the other peptides formed a more or less similar number of H-bonds with the LecA protein (Figure 2e).

**Figure 2.**
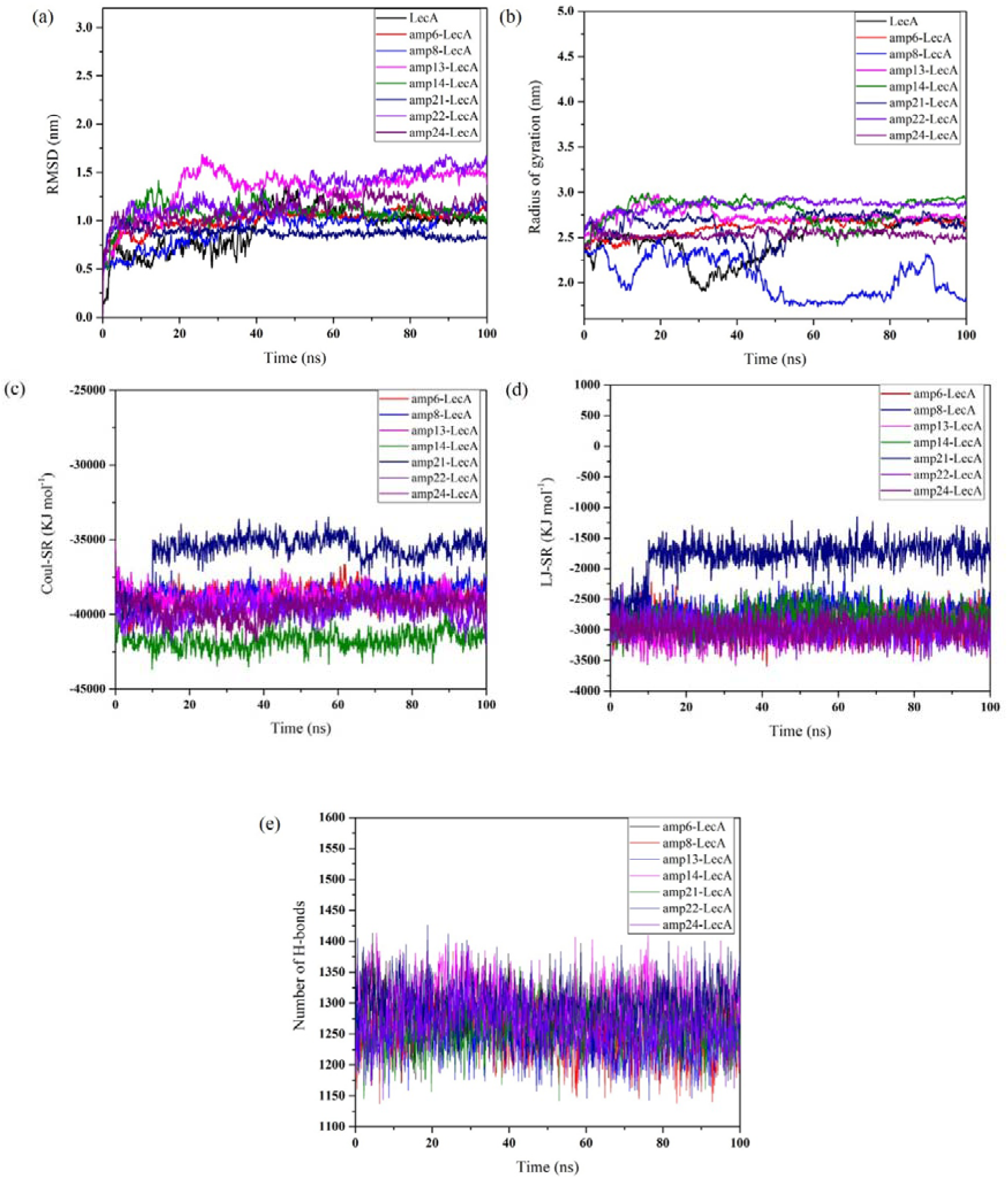
**(a)** RMSD profile of LecA-peptide complexes **(b)** Rg profile of LecA-peptide complexes, **(c)** Coul-SR profile for LecA-peptide complexes, **(d)** LJ-SR profile for LecA-peptide complexes, **(e)** H-Bonds for LecA-peptide complexes

Further, amp24-LecA and amp21- complexes showed a significant number of conformational states in the blue region or the lower energy state, indicating the stability of these complexes (Figure 3). However, based on overall MD simulation analysis of LecA-peptide complexes, amp24 and amp21 seem to be binding most stably with LecA.

**Figure 3.**
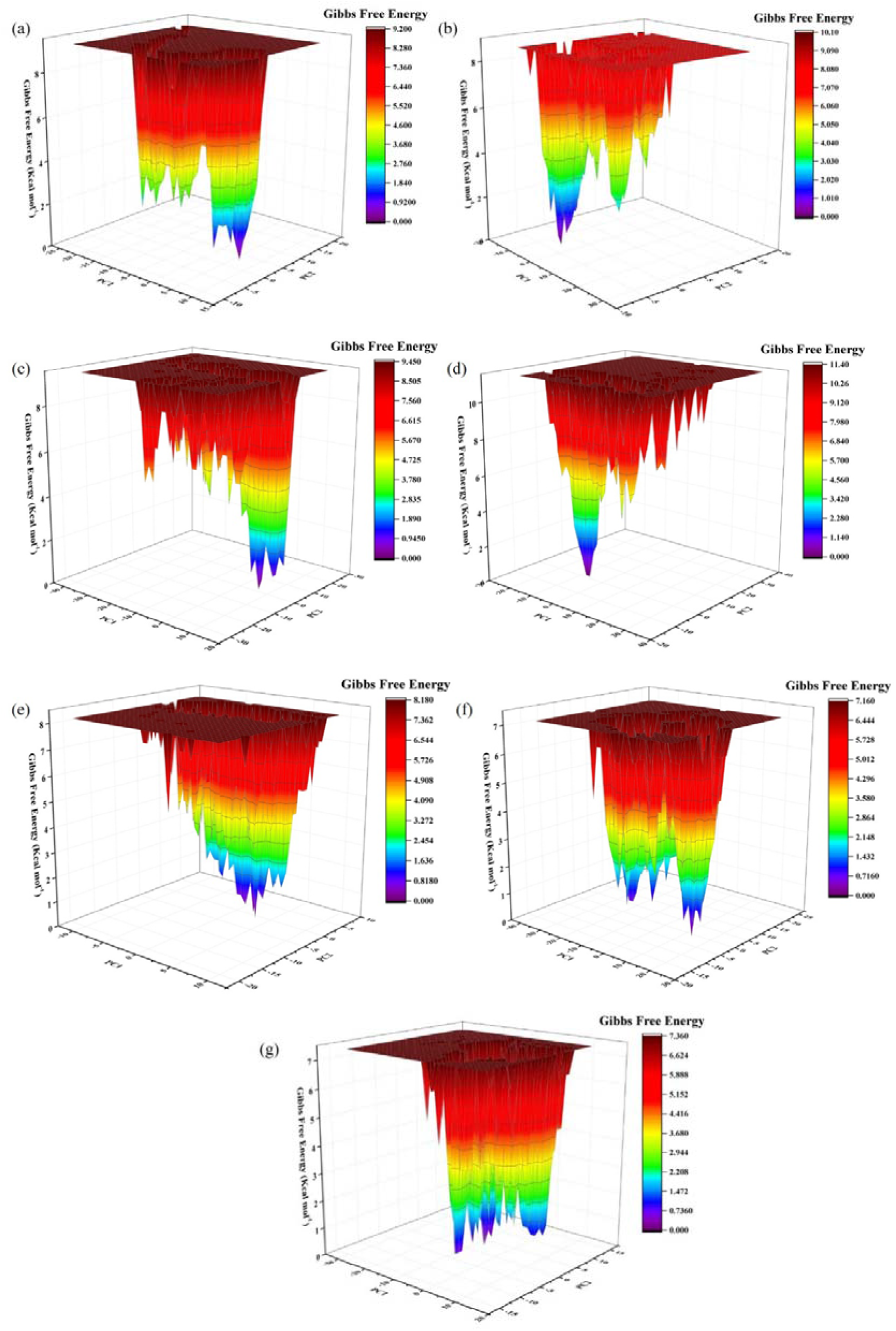
FEL profile of (a) amp6-LecA **(b)** amp8-LecA **(c)** amp13-LecA **(d)** amp14-LecA **(e)** amp21-LecA **(f)** amp22-LecA **(g)** amp24-LecA

#### 3.5.2. Analyzing the dynamic stability and interaction profiling of LecB–peptide complexes

As compared to other LecB-peptide complexes, amp21-LecB complex and amp6-LecB showed lower RMSD values. This indicated that both amp21 and amp6 maintained a stable and comparatively consistent binding at the interaction site in LecB protein (Figure 4a). The complex amp6-LecB showed the least fluctuations in Rg; significant results were not obtained in Rg plots of other LecB-peptide complexes(Figure 4b). amp14-LecB and amp22-LecB showed the lowest Coul-SR values among all the LecB-peptide complexes (Figure 4c). amp14-LecB and amp13-LecB showed the lowest LJ-SR values (Figure 4d). Among all the AMPs, amp14 formed the maximum number of H-bonds with LecB (Figure 4e).

**Figure 4.**
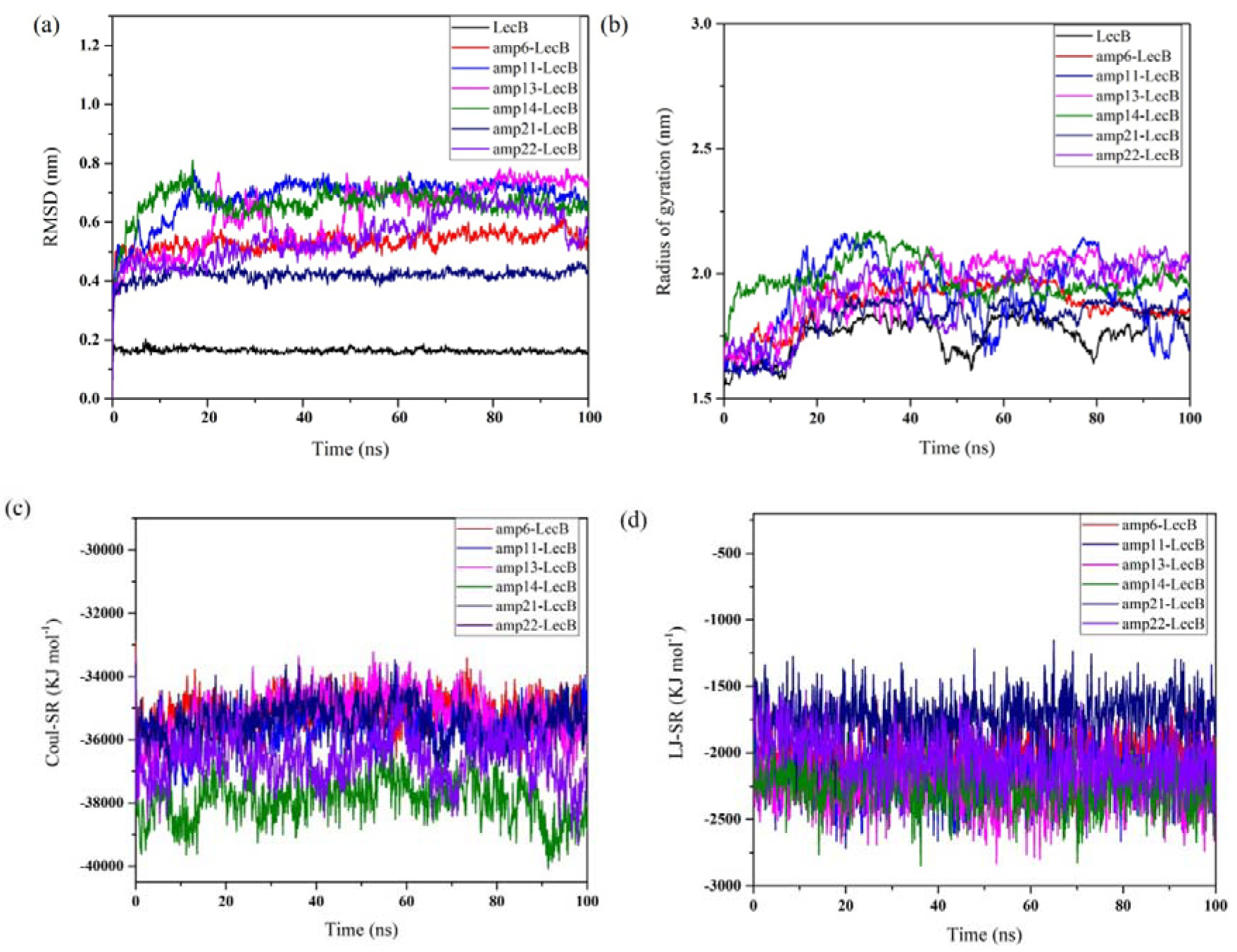
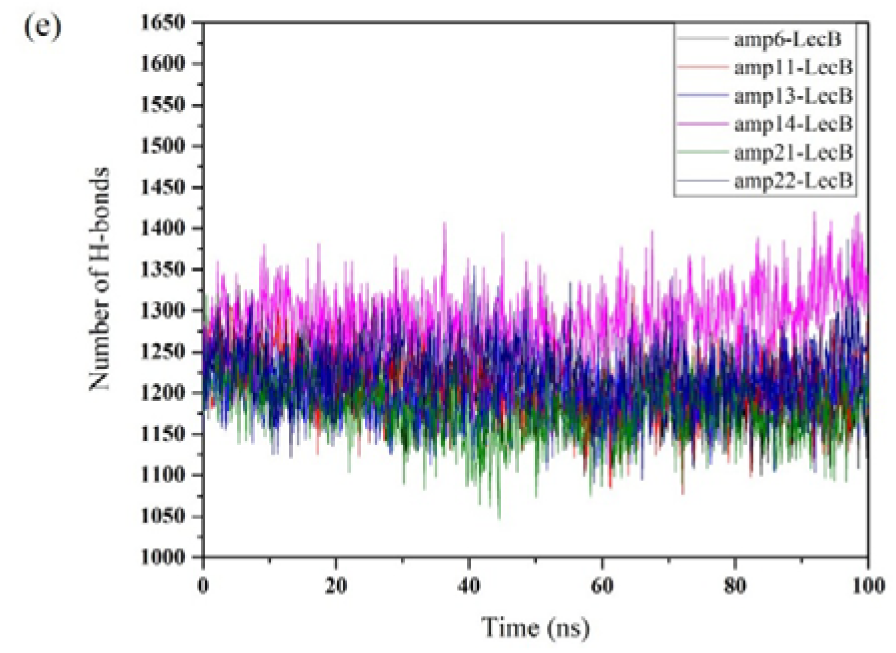
**(a)** RMSD profile of LecB-peptide complexes, **(b)** Rg profile of LecB-peptide complexes, **(c)** Coul-SR profile for LecB-peptide complexes, **(d)** LJ-SR profile for LecB-peptide complexes, **(e)** H-Bonds for LecB-peptide complexes

The amp13-LecB, amp22-LecB, and amp21-LecB complexes showed a significant number of conformational states in the blue region or the lower energy state, indicating the stability of these complexes (Figure 5). However, amp13 and amp22 showed very large principal component analysis (PCA) dimensionality, indicating unstable conformations or scattered states of complexes (Figure 5). Only amp21 and amp6 bound LecB complexes show low PCA dimensions (Figure 5). Thus, based on overall MD simulation analysis of LecB-peptide complexes, amp6 and amp21 seem to be binding most stably with LecB.

**Figure 5.**
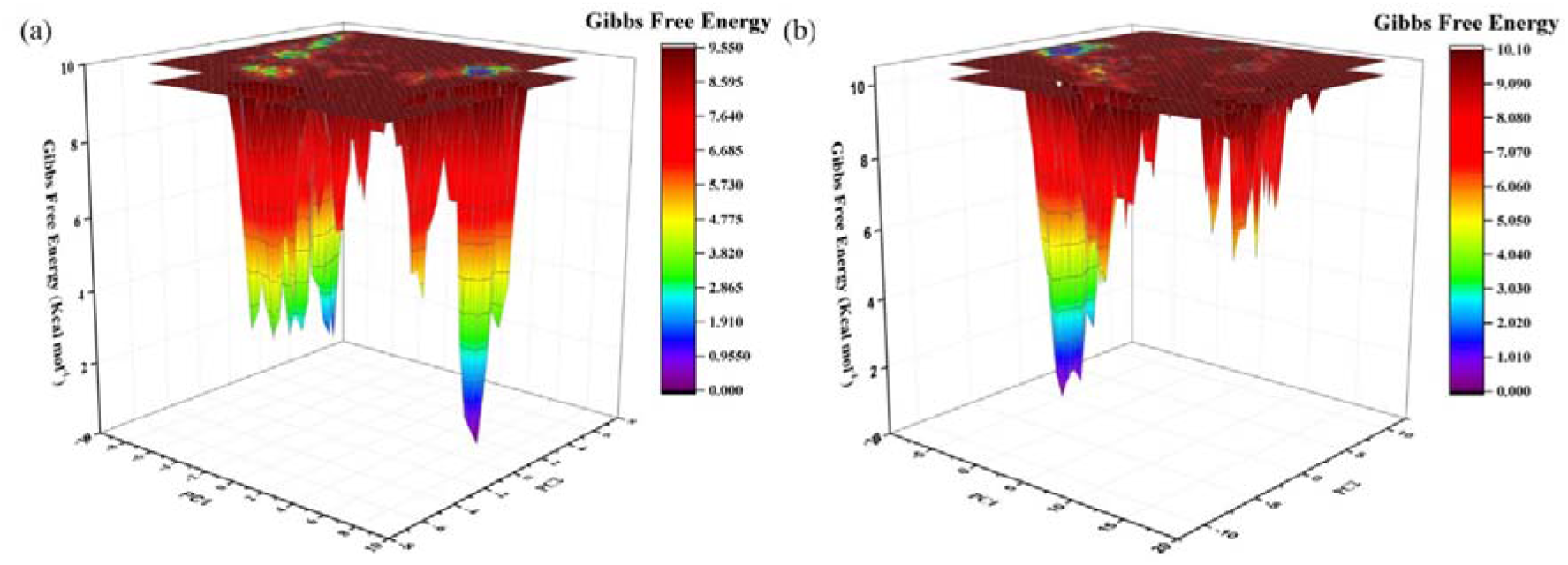
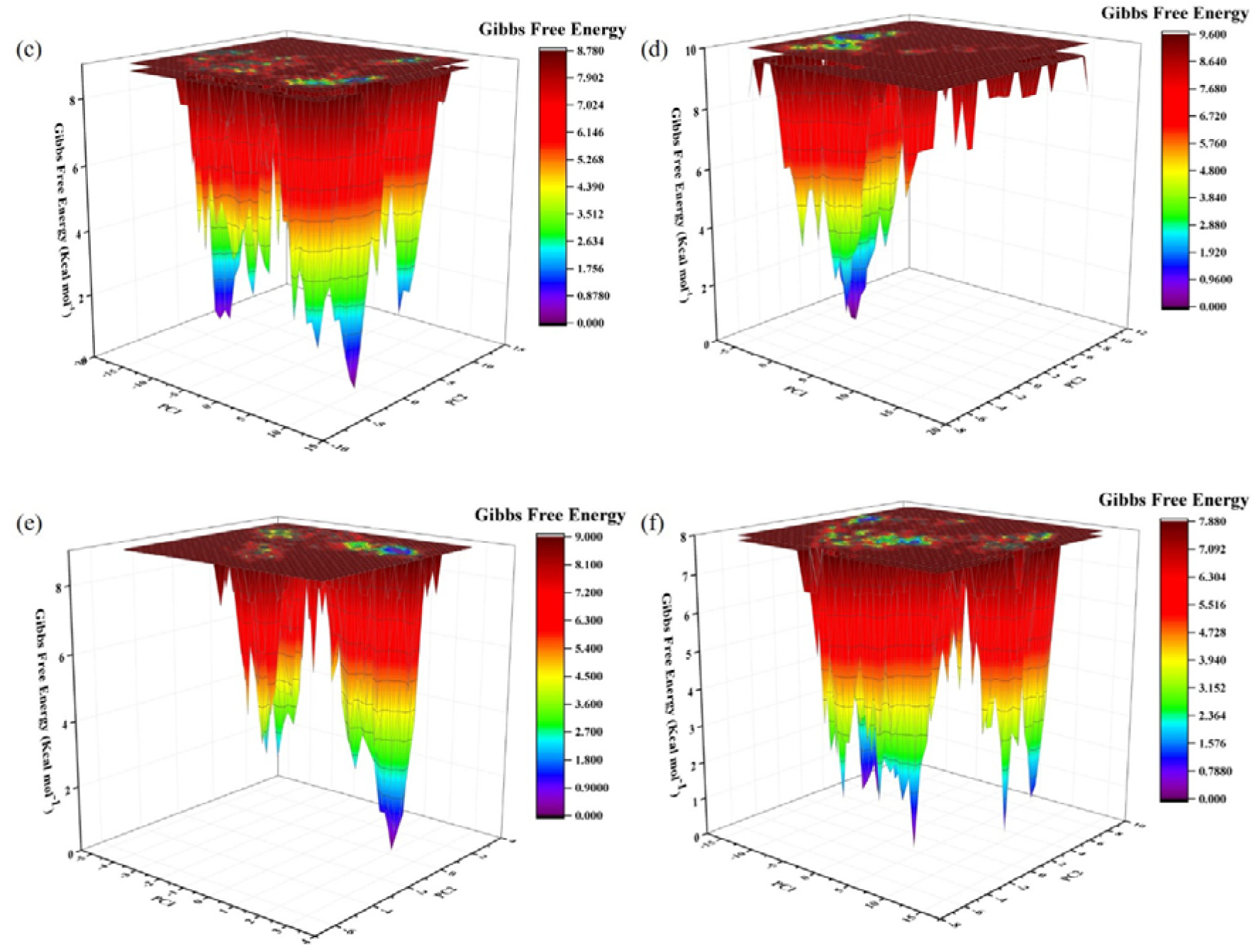
**(a)** FEL profile of amp6-LecB **(b)** amp11-LecB **(c)** amp13-LecB **(d)** amp14-LecB **(e)** amp21-LecB **(f)** amp22-LecB

Based on MD simulation analysis results, it was inferred that amp21 and amp24 showed the best binding affinity with LecA protein, as (i) amp21-LecA complex showed minimum RMSD with a good FEL plot, (ii) amp24-LecA complex showed low and constant Rg with a good FEL plot. amp6 and amp21 showed the best binding affinity with LecB protein, as (i) amp21-LecB complex showed minimum RMSD with a good FEL plot, (ii) amp6-LecB complex showed the least fluctuations in Rg with low RMSD and LJ-SR.

### 3.6. Selection of topmost antibiofilm peptides

As is evident from the MD simulation analysis results, LecA protein formed the most stable complexes with amp21 and amp24, while LecB formed the most stable complexes with amp6 and amp21. These *in silico* findings guided the selection of amp6, amp21, and amp24 for subsequent experimental validation. The above peptides were sent for synthesis to Peptide Purchase Committee, Elabscience, USA, having more than 95% purity. From the chosen peptides, amp21 has been mentioned as AMP218 whereas amp24 has been mentioned as AMP2410 by Anand et al. [Anand et al., 2025].

### 3.7. Comparative study of the residue-level binding of lead AMPs with known raffinose binding sites in LecA protein

amp21 and amp24 were identified as the strongest binders to LecA protein based on MD simulation results. Therefore, their binding residues were analyzed to assess whether they overlap with the known raffinose-binding sites of LecA. Raffinose has been previously documented to inhibit biofilm formation in *P. aeruginosa* by inhibiting the functioning of LecA protein [Kim et al., 2016]

In order to understand the binding of raffinose (antibiofilm agent) to LecA, Kim and coworkers performed *in silico* docking studies between raffinose and X-ray structure of LecA in complex with melibiose. Most of the docked raffinose poses showed common binding patterns in which the galactose moiety of raffinose was projected toward the calcium ion of the LecA active site, while the glucose and fructose moieties of raffinose were oriented toward the surface and the loop region of LecA. The galactose moiety of the raffinose interacted strongly with Tyr 36, His 50, Pro 51, Asp 100, Val 101, and Thr 104, the glucose interacted by hydrogen-bonding with Gln 53, the fructose interacted by hydrogen-bonding with Gly 37 and Asn 107 [Kim et al., 2016].

Since, only amp21 and amp24 were top candidates showing good binding to lecA, we performed a comparative analysis with raffinose binding sites of LecA, and found interesting results. Both amp21 and amp24 were found to interact with all three raffinose binding sites, that is, binding site of raffinose, fructose as well as glucose moiety. The results for same can be visualized in Figure 6, suggesting the possible role of AMPs in biofilm inhibition in a manner similar to raffinose.

**Figure 6.**
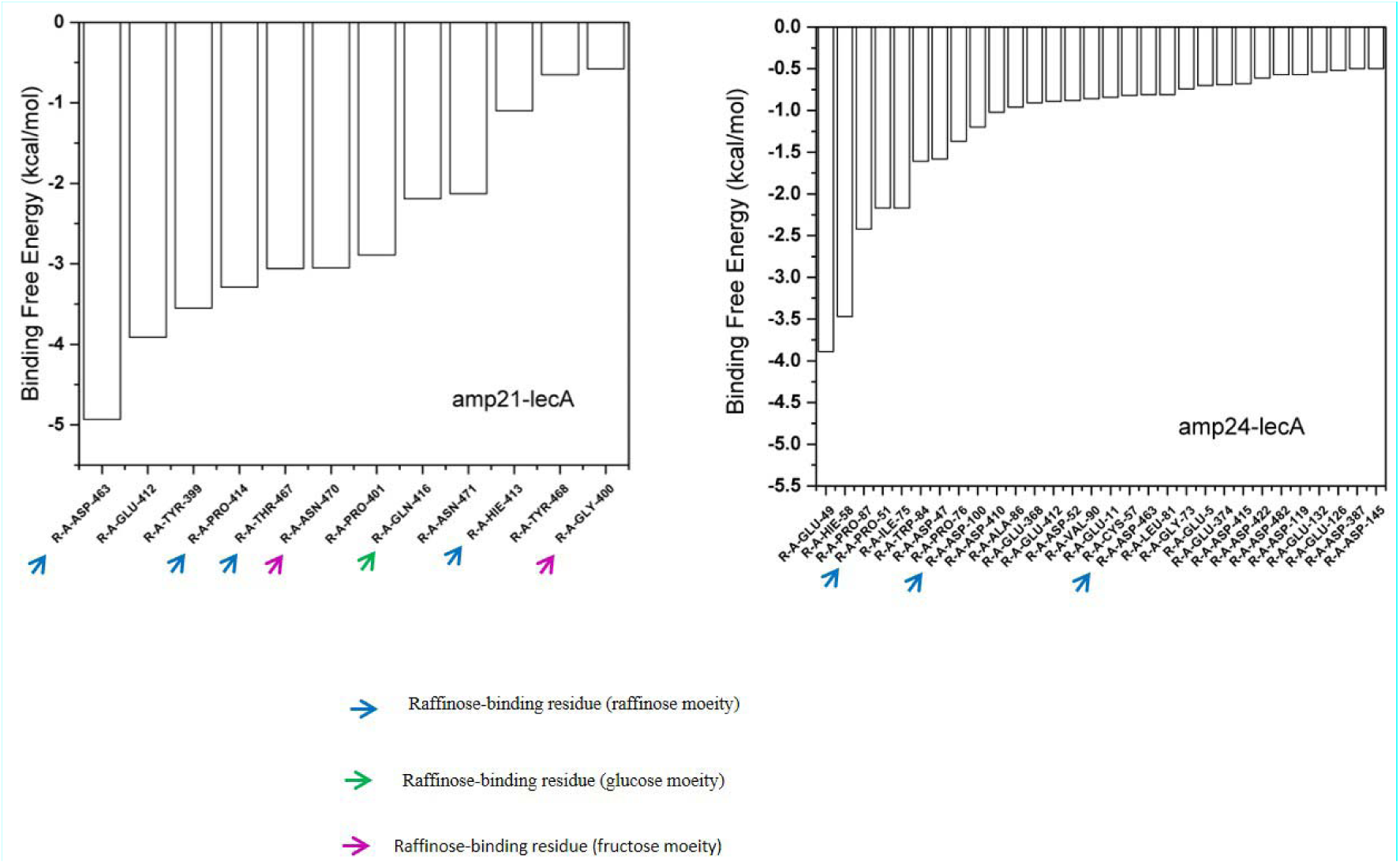
Binding of amp21 and amp24 with raffinose-binding residues of LecA

### 3.8. Evaluation of the hemocompatibility of lead AMPs

Before studying the antibacterial activity of amp6, amp21, and amp24, their hemocompatibility was assessed using human RBCs. Upto a concentration of 100 µg mL^-1^, all three AMPs induced minimal hemolysis and affected a nominal fraction of the total RBC population (Figure 7). At 200 µg mL^-1^, amp6 and amp21 caused 2.85% and 0.99% hemolysis, respectively (Figure 7). However, amp24 exhibited comparatively higher hemolytic activity at 200 µg mL^-1^ and resulted in 7.5% hemolysis. Overall, these findings demonstrated that amp6, amp21, and amp24 were hemocompatible over a broad range of concentration.

**Figure 7.**
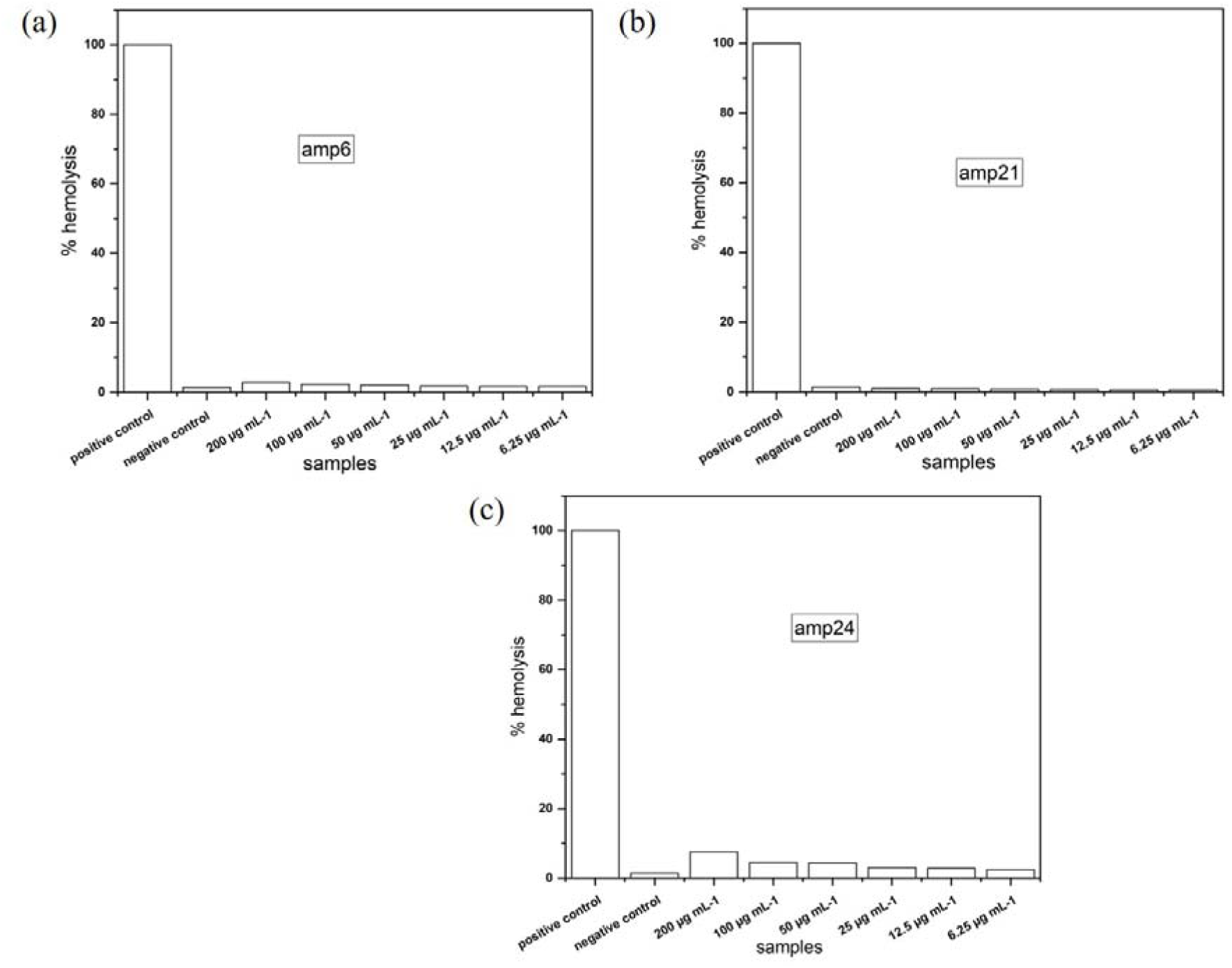
Hemolysis assay of **(a)** amp6 **(b)** amp21 **(c)** amp24

### 3.9. Determining the cytotoxicity of lead AMPs against mammalian cell lines

Since no hemolytic activity was observed up to 200 µg mL□¹ of AMPs, cytotoxicity assay was performed within a concentration range of 6.25- 100 µg mL□¹ to avoid unnecessary exposure of the cells to high concentrations of AMPs. Out of the 3 AMPs, amp21 was found to be non-toxic to HCT116 cells across the entire tested concentration range (Figure 8a). Even at 100 µg mL^-1^, amp21-treated cells exhibited a viability of 105.6% relative to the untreated control (100%). amp6 did not affected the viability of HCT116 cells up to 50 µg mL^-1^. However, a marginal decrease in the viability was observed in HCT116 cells at 100 µg mL^-1^ of amp6 (Figure 8b). amp24 showed no significant effect on the cell viability up to 25 µg mL^-1^. However, the viability of HCT116 cells decreased by nearly 20% at 50 µg mL-1 and by 28% at 100 µg mL^-1^ of amp24 (Figure 8c).

**Figure 8.**
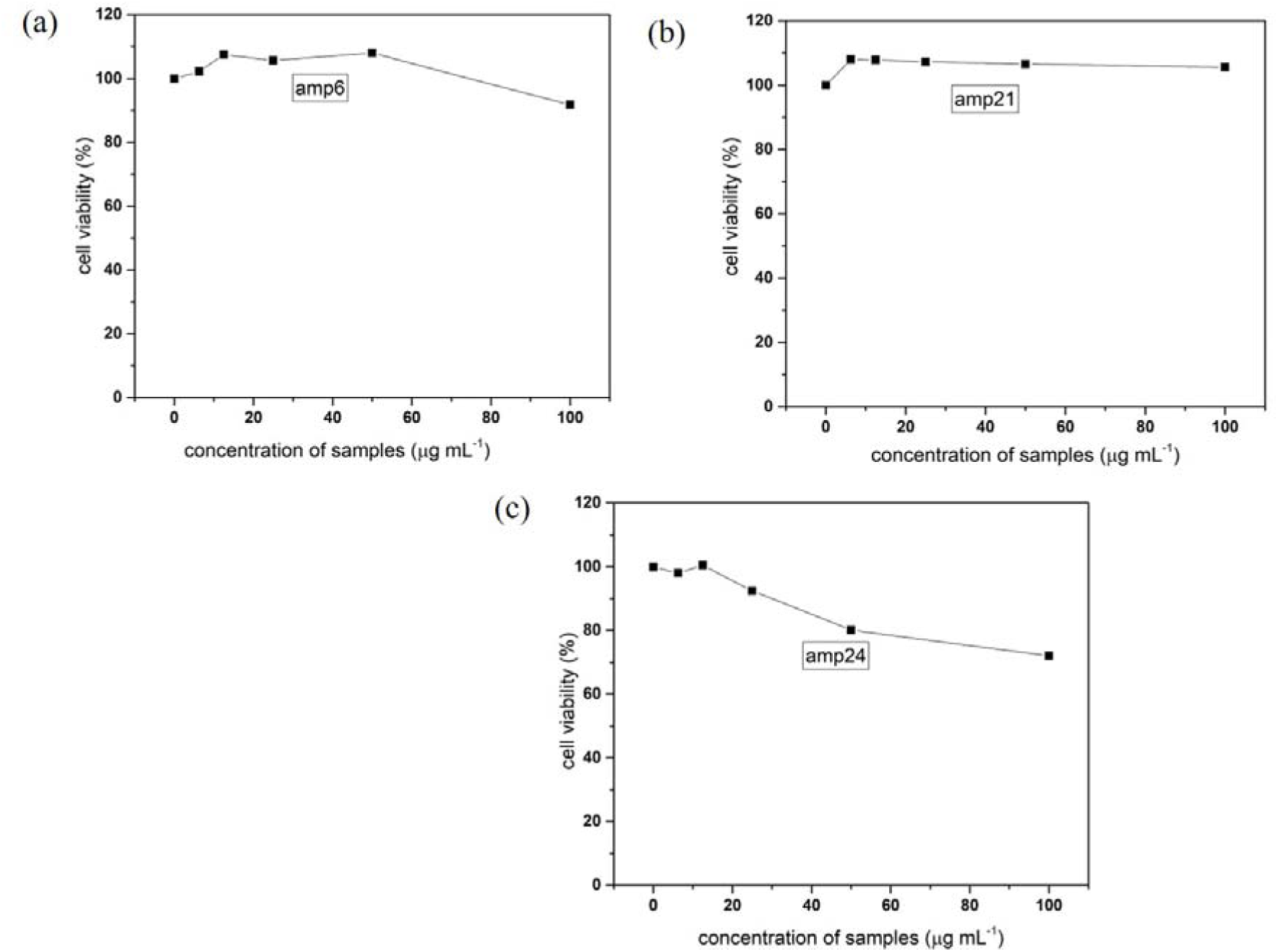
Cytotoxicity assay of **(a)** amp6 **(b)** amp21 **(c)** amp24

### 3.10. Studying the dose-dependent antibacterial activity of lead AMPs against *P. aeruginosa* through CFU assay

Following confirmation of hemocompatibility and acceptable cytotoxicity profiles, the antibacterial efficacy of the selected AMPs was evaluated using the colony-forming unit (CFU) counting assay. amp21 exhibited the strongest antibacterial activity against *P. aeruginosa*, reducing CFU counts by approximately 60% and 84% CFU mL□¹ at concentrations of 50 µg mL□¹ and 100 µg mL□¹, respectively (Figure 9). amp6 showed a comparatively moderate antibacterial activity with CFU reductions of nearly 38% and 64%, respectively, at 50 µg mL□¹ and 100 µg mL□¹, respectively (Figure 9). Interestingly, amp24 showed comparable antibacterial efficacy at both concentrations, resulting in an approximately 54% reduction in CFU counts (Figure 9).

**Figure 9.**
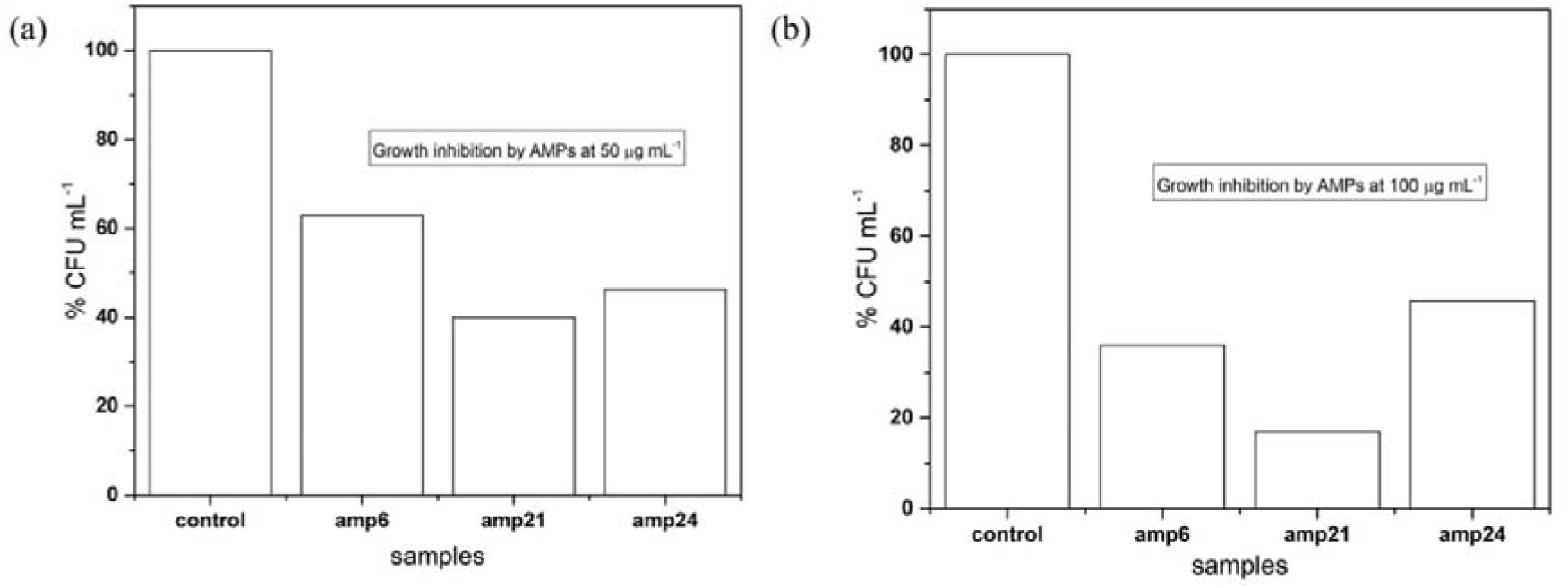
CFU estimation of planktonic cells of *P. aeruginosa* on treatment with amp6,amp21, and amp24 at **(a)** 50 µg mL^-1^ and **(b)** 100 µg mL^-1^ concentration.

### 3.11. Permeabilization of bacterial cell membranes by lead AMPs assessed through PI uptake

To elucidate the underlying mechanism for the observed antibacterial activity of the lead AMPs, membrane permeabilization of *P. aeruginosa* cells was evaluated through the PI uptake experiment. On comparison with the untreated cells, *P. aeruginosa* cells treated with 12.5 µg mL^-1^ and 100 µg mL^-1^ of the lead AMPs resulted in a marked increase in PI fluorescence (Figure 10). The above increase in PI fluorescence in AMP-treated cells indicated that all of the above AMPs compromised the integrity of the bacterial cell membrane. The above phenomenon led to an increase in PI uptake by the bacterial cells and subsequent increased binding to intracellular DNA.

**Figure 10.**
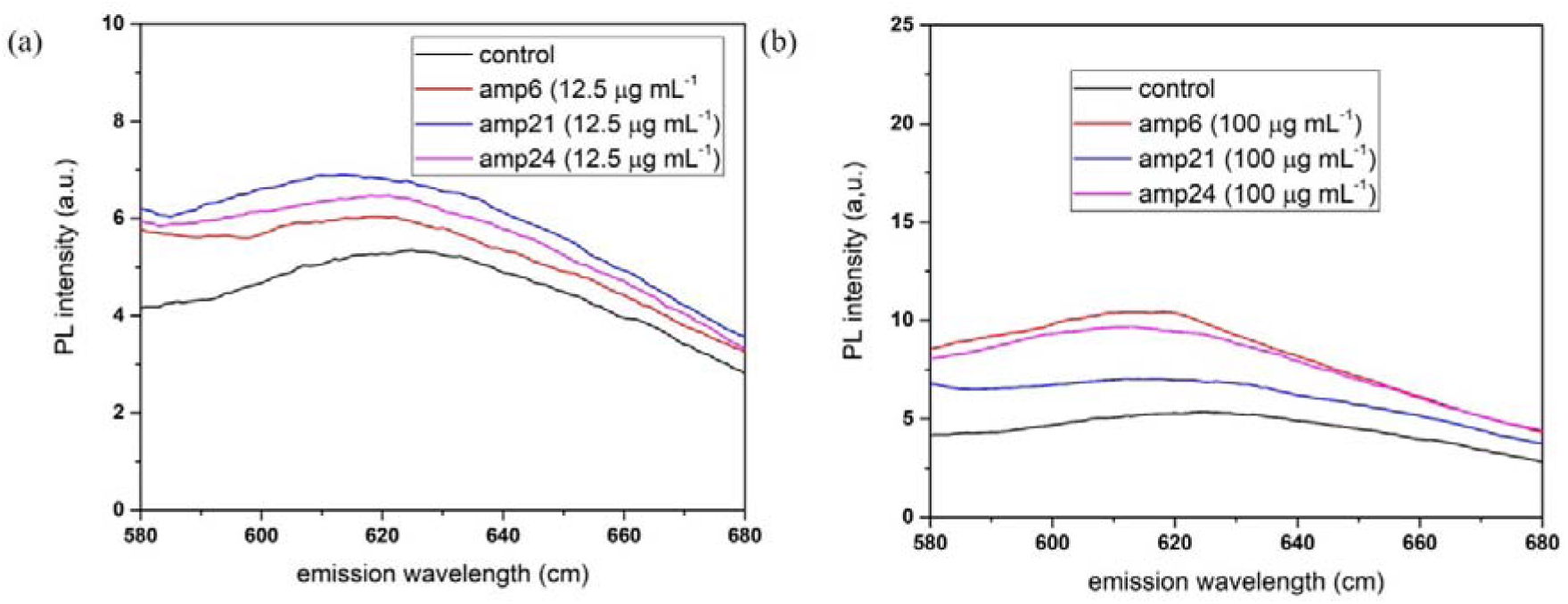
Relative fluorescence of PI uptake to assess membrane permeability of *P. aeruginosa* on treatment with amp6, amp21, and amp24 at **(a)** 12.5 µg mL^-1,^ **(b)** and 100 µg mL^-1^ concentration. Herein, the untreated *P. aeruginosa* cells were used as the control

### 3.12. Assessment of antibiofilm assessment of AMPs

The antibiofilm potential of the lead AMPs against *P. aeruginosa* was assessed through biofilm inhibition and biofilm disruption assays after evaluating their antibacterial efficacy:

#### 3.12.1. Quantification of *P. aeruginosa* biofilm inhibition by AMPs

It was found that treatment of *P. aeruginosa* cells with 50 µg mL^-1^ of amp6 and amp21 before biofilm formation significantly inhibited biofilm development, reduced the biofilm biomass to 63.4% and 67.9% of the untreated control, respectively (Figure 11). However, amp24 showed comparatively lower biofilm inhibition, where the biofilm mass was reduced to 80.9% relative to the untreated control under identical conditions (Figure 11).

**Figure 11.**
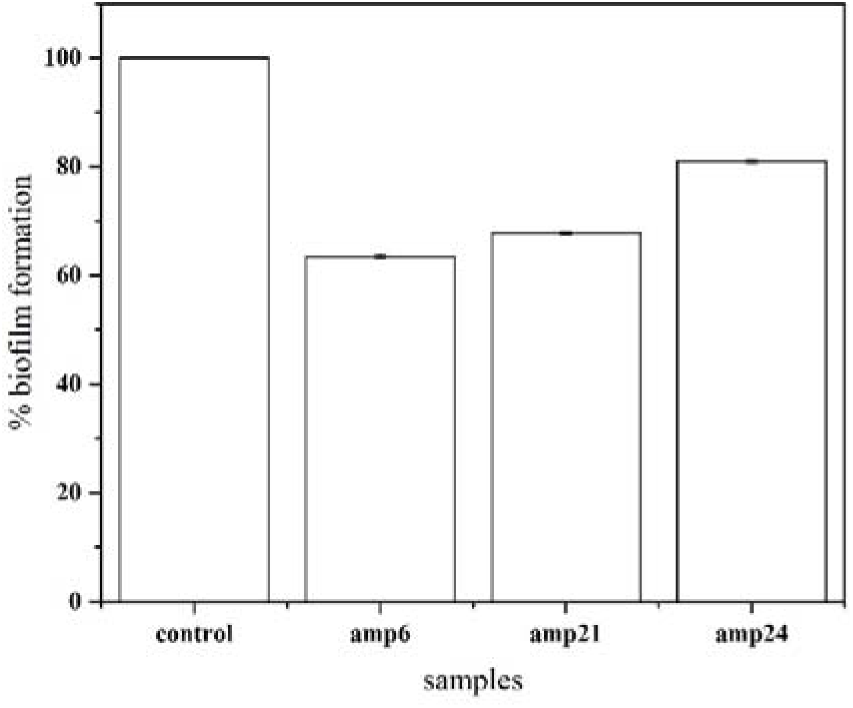
Quantitative estimation of *P. aeruginosa* biofilm inhibition by 50 µg mL^-1^ of amp6, amp21, and amp24. Herein, the untreated biofilm was used as the control.

#### 3.12.2. Quantification of *P. aeruginosa* biofilm disruption by AMPs

Treatment of preformed *P. aeruginosa* biofilms with 50 µg mL^-1^ of amp6 and amp21 resulted in substantial biofilm disruption, reducing the biofilm biomass to 68.18% and 63.6% of the untreated control, respectively (Figure 12). amp24 demonstrated relatively lower biofilm-disruption efficacy, where the biofilm mass was reduced to 75.75% relative to the control under the same conditions (Figure 12).

**Figure 12.**
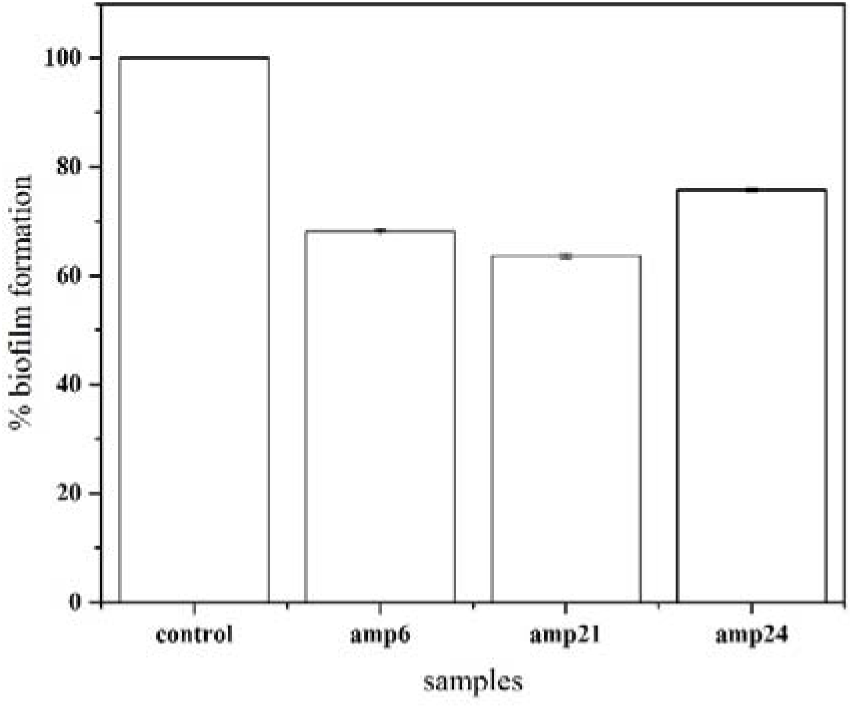
Quantitative estimation of *P. aeruginosa* biofilm disruption by 50 µg mL^-1^ of amp6,amp21, and amp24.

### 3.13. Studying the AMPs induced morphological changes in *P. aeruginosa* biofilms through SEM imaging

To visually corroborate the biofilm disruption observed in section 3.12.2, we subsequently performed scanning electron microscopy (SEM) analysis. Due to methodological constraints, economic considerations, and observation of similar functional outcomes for all three AMPs in the PI fluorescence assay, SEM analysis was performed only for amp6, which served as a representative AMP. SEM analyses revealed that when compared to the untreated (control) biofilm, *P. aeruginosa* biofilms treated with amp6 showed pronounced structural disruption (Figure 13b). The untreated biofilms appeared largely intact, dense, and well-organised (Figure 13a). On the other hand, a marked reduction was observed in the bacterial cell density of amp6-treated biofilms (Figure 13b). Furthermore, treatment with amp6 not only disrupted the EPS matrix but cause prominent disruption to the bacterial cell membrane (Figure 13b). Similar effects are anticipated as both peptides, like amp6, compromise the integrity of the bacterial cell membrane and show comparable biofilm inhibition and disruptive activity.

**Figure 13.**
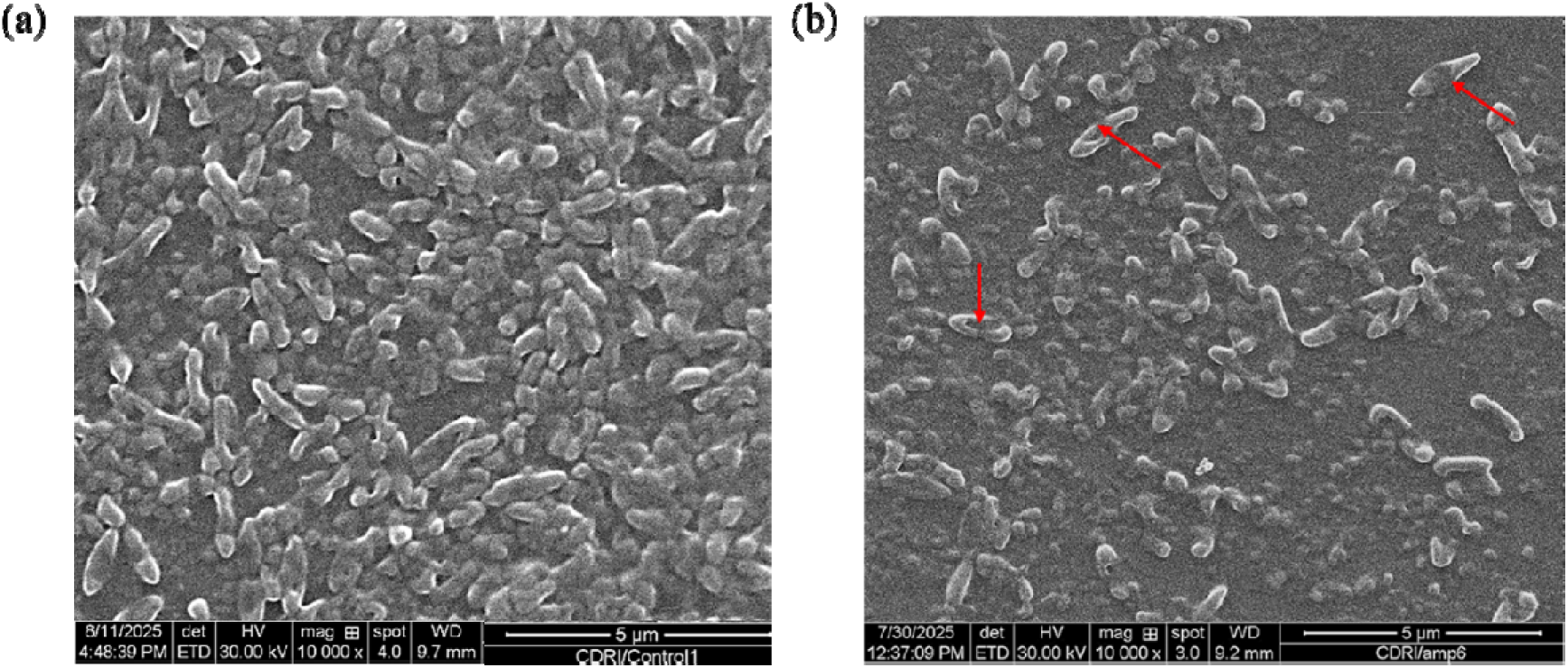
SEM image of **(a)** untreated *P. aeruginosa* biofilm, **(b)** *P. aeruginosa* biofilm treated with amp6. The damaged bacterial cells are shown by the red arrow

### 3.14. Mechanistic insights into biofilm disruption mediated by AMPs

Following the quantitative assessment of biofilm disruption and its morphological validation by SEM imaging, we elucidated the underlying mechanism of antibiofilm activity of the selected AMPs. Upon treatment of mature *P. aeruginosa* biofilms with amp6, amp21, and amp24, it was observed that there was a marked increase in the number of bacterial cells migrating from the biofilm pellicle into the underlying media (Figure 14a). The cells released from the biofilm matrix were subsequently quantified through the CFU assay. A substantially higher number of bacterial colonies were detected on Mueller–Hinton agar plates inoculated with AMP-treated cultures (Figure 14b). The above observations suggested that the above three AMPs exerted their antibiofilm activity by disrupting the integrity of the biofilm EPS matrix, which eventually led to the migration of underlying bacteria to the planktonic state. The above phenomenon may be attributed to the interaction of AMPs with lectin proteins that subsequently disturbed the integrity of the biofilm EPS matrix.

**Figure 14.**
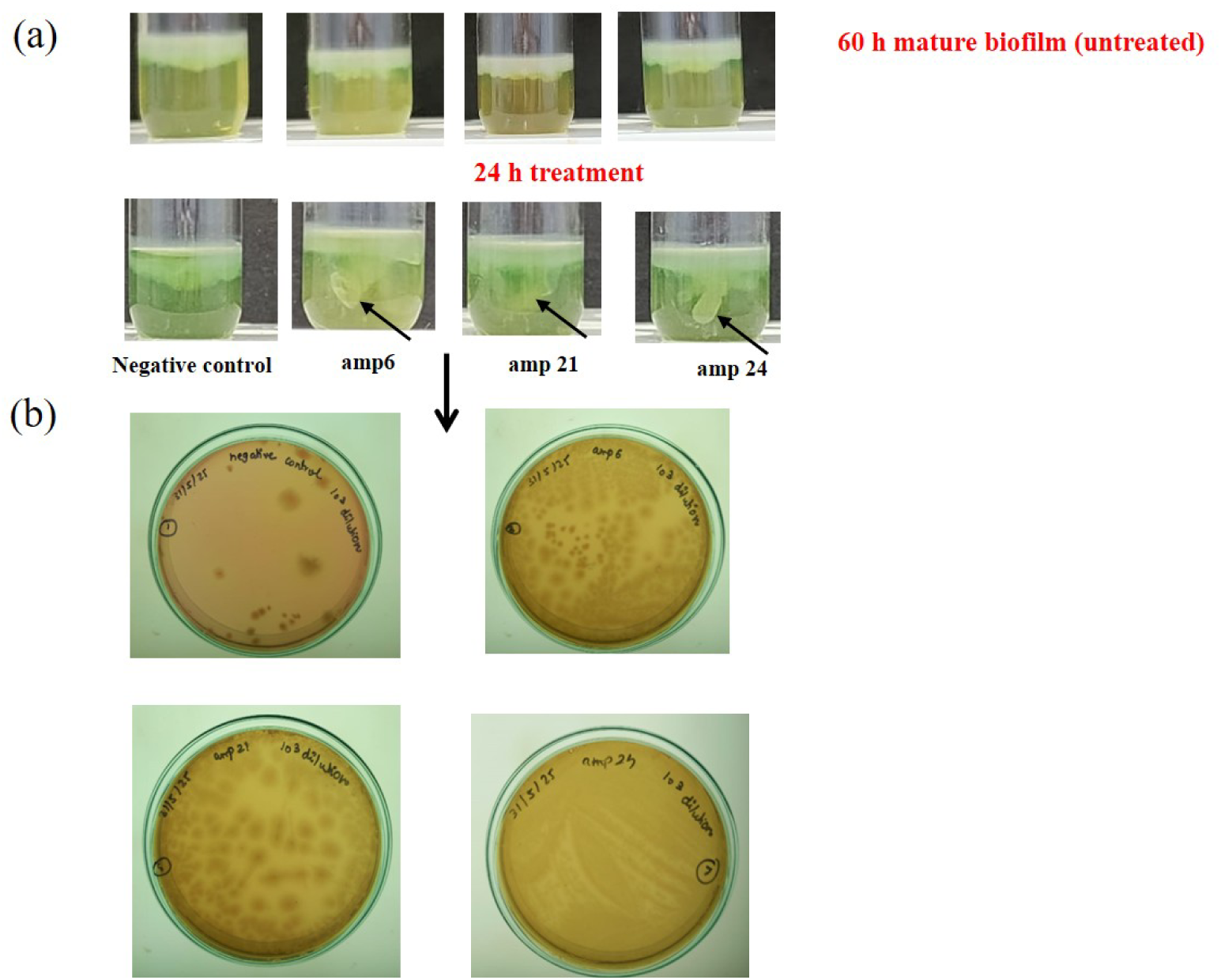
**(a)** Movement of bacterial cell from biofilm pellicle to the surrounding media showing transition from biofilm to the free planktonic cells **(b)** CFU assay to visualize the planktonic cell load in the media surrounding the biofilm on treatment with AMPs. Herein, the untreated *P. aeruginosa* cells were used as the control

### 3.15. Designing ultrashort peptides (USPs) from lead AMPs targeting the LecA protein

In order to address the limitations of the full length parent AMPs such as high cost of synthesis, hemocompatibility and cytotoxicity issues at higher concentrations, we designed the ultrashort derivatives of the parent peptides that showed best binding affinity to LecA protein. From the MD simulation studies, it was evident that amp21 and amp24 showed best binding affinity with LecA protein. Therefore, we derived USPs from amp21 and amp24 and investigated their binding efficacy with LecA protein through molecular docking, MM-GBSA and MD simulation studies.

#### 3.15.1. Analysing the interaction pattern between AMP residues and carbohydrate binding regions of the LecA protein

Using the RING web server, we investigated the patterns of interaction between the residues of amp21 and amp24 with the carbohydrate binding regions of the LecA protein, more especially, those connected to the galactose, glucose, and fructose moieties of raffinose. The lysine at position 14 of amp21 consistently formed both hydrogen-bond and ionic interactions with the galactose-binding residue of raffinose, according to the analysis (Figure 15(a-b)). The interaction frequency surpassed 50% in the majority of the trajectory’s frames, suggesting a consistent and recurrent contact. Apart from this residue, it was also discovered that arginine at position 22 engages in ionic interactions with the same galactose moiety.

**Figure 15.**
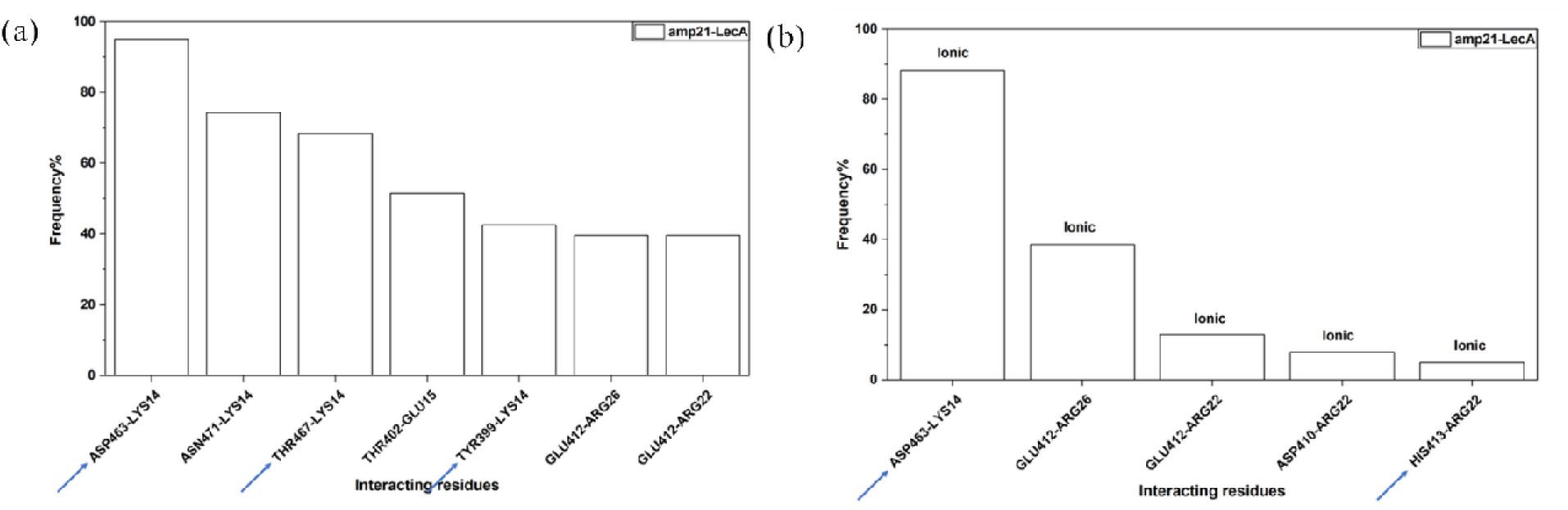
**(a)** Frequency of H-bond interaction between residues of amp21 with LecA protein during MD simulation **(b)** Frequency of ionic interaction between residues of amp21 with LecA protein during MD simulation

For amp24, a comparable pattern was noted (Figure 16). While the lysine at position 4 formed hydrogen bonds with the glucose-interacting residue, the lysine residues at positions 1 and 3 frequently engaged in hydrogen-bond interactions with the galactose-associated residues of raffinose. This analysis helped us identify which specific residues in the peptide were actually engaging with the various raffinose-related binding sites. The design of the ultra-short peptides began with these crucial interaction residues.

**Figure 16.**
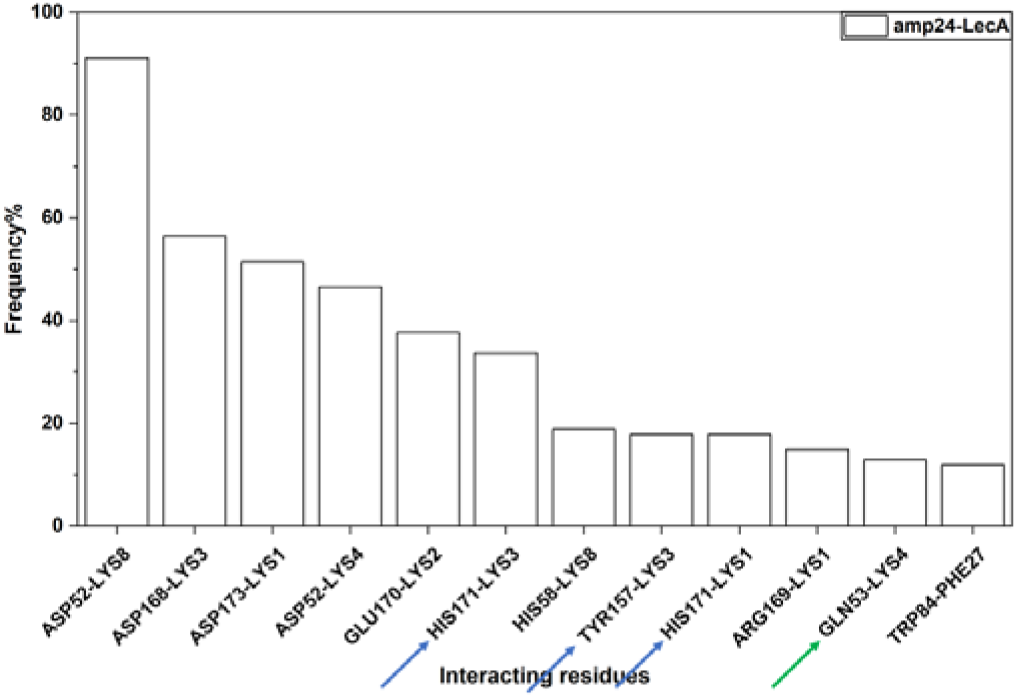
Frequency of H-bond interaction between residues of amp24 with LecA protein during MD simulation

#### 3.15.2. Designing USPs from amp21 and amp24

Using the insights from the RING analysis, we put together six different USP sequences for each of the parent peptides by designing them manually. The design approach for amp21 was quite simple; we only chose the original sequence segments that contained residues that were directly involved in interactions with the LecA carbohydrate-binding regions. But for amp24, things were different. Here, only the N-terminal lysine residue participated in the interaction, and the peptide sequence itself contained repeated stretches of lysine and phenylalanine. Because of this repetition, it was not possible to generate six distinct sequence variants by following the same approach used for amp21. To address this, we selected a series of short peptide fragments from the central region of the parent amp24 sequence that naturally include the interacting lysine while preserving the original residue order, allowing us to produce a meaningful set of USPs. The final names, sequences, and lengths of the USPs derived from amp21 and amp24 are presented in Tables 4 and 5, respectively.

**Table 4.** USPs designed from amp21.

| S. no. | Name | Sequence | Length |
| --- | --- | --- | --- |
| 1. | amp21.1 | QKEKEN | 6 |
| 2. | amp21.2 | EKENE | 5 |
| 3. | amp21.3 | KEKENE | 6 |
| 4. | amp21.4 | LTRRRL | 6 |
| 5. | amp21.5 | RRRLSR | 6 |
| 6. | amp21.6 | TRRRLSR | 7 |

**Table 5.** USPs designed from amp24.

| S. no. | Name | Sequence | Length |
| --- | --- | --- | --- |
| 1. | amp24.1 | KKKKR | 5 |
| 2. | amp24.2 | KKRKKK | 6 |
| 3. | amp24.3 | KRKKKKK | 7 |
| 4. | amp24.4 | KKRVFFF | 7 |
| 5. | amp24.5 | KKKKRVF | 7 |
| 6. | amp24.6 | KRVFFF | 6 |

#### 3.15.3. Studying the binding affinity of the USPs with LecA protein through molecular docking and MM/GBSA analysis

Using the HawkDock server, we performed the docking and MMGBSA analysis for the USPs derived from amp21 and amp24. The docking and MMGBSA scores are listed in Table 6 and 7 for derivatives of amp21 and in Table 8 and 9 for derivatives of amp24.

**Table 6.** Docking score of USPs derived from amp21, which is sorted in ascending order.

| S. no. | USPs derived from amp21 | Docking Score |
| --- | --- | --- |
| 1. | amp21.5 | -3268.73 |
| 2. | amp21.6 | -3248.67 |
| 3. | amp21.4 | -3207.58 |
| 4. | amp21.1 | -2247.64 |
| 5. | amp21.3 | -2148.64 |
| 6. | amp21.2 | -2051.35 |

**Table 7.** MMGBSA score of USPs derived from amp21, which is sorted in ascending order.

| S. no. | USPs derived from amp21 | MMGBSA Score (kcal mol<sup>-1</sup>) |
| --- | --- | --- |
| 1. | amp21.6 | -66.99 |
| 2. | amp21.4 | -51.11 |
| 3. | amp21.5 | -50.27 |
| 4. | amp21.1 | -29.1 |
| 5. | amp21.2 | -28.52 |
| 6. | amp21.3 | -25.86 |

**Table 8.**
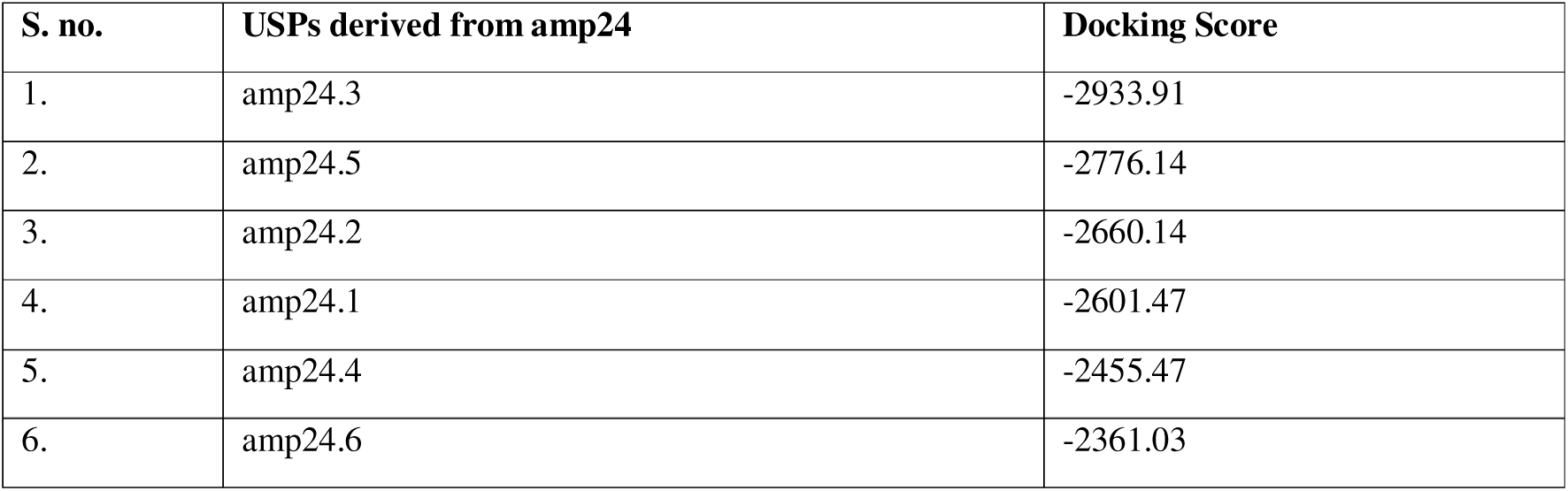
Docking score of USPs derived from amp24, which is sorted in ascending order.

**Table 9.** Docking score of USPs derived from amp24, which is sorted in ascending order.

| S. no. | USPs derived from amp24 | MMGBSA Score (kcal/mol) |
| --- | --- | --- |
| 1. | amp24.2 | -66.99 |
| 2. | amp24.5 | -51.11 |
| 3. | amp24.6 | -50.27 |
| 4. | amp24.4 | -29.1 |
| 5. | amp24.1 | -28.52 |
| 6. | amp24.3 | -25.86 |

#### 3.15.4. Analyzing the 2D interaction between USPs and LecA protein

From each set of amp21- and amp24-derived USPs, we selected the top-performing USPs for 2D interaction analysis. The selection was based on a combination of docking scores and MM/GBSA values. For the amp21 group, the USPs amp21.4, amp21.5, and amp21.6 were chosen, while from the amp24 group, amp24.2, amp24.3, amp24.5, and amp24.6 were shortlisted. The interaction patterns of these peptides with LecA were examined using LigPlot.

The 2D interaction results for the amp21 derivatives (Figure 17) showed that all three selected USPs formed favorable contacts with residues associated with the raffinose-binding regions of LecA, including those linked to the galactose, glucose, and fructose moieties. In the case of the amp24 derivatives (Figure 18), three out of the four USPs, amp24.2, amp24.3, and amp24.6, displayed consistent and meaningful interactions with the raffinose-binding residues, while one peptide showed weaker involvement. Based on these findings, a total of six USPs, three from amp21 and three from amp24, were selected for subsequent MD simulation.

**Figure 17.**
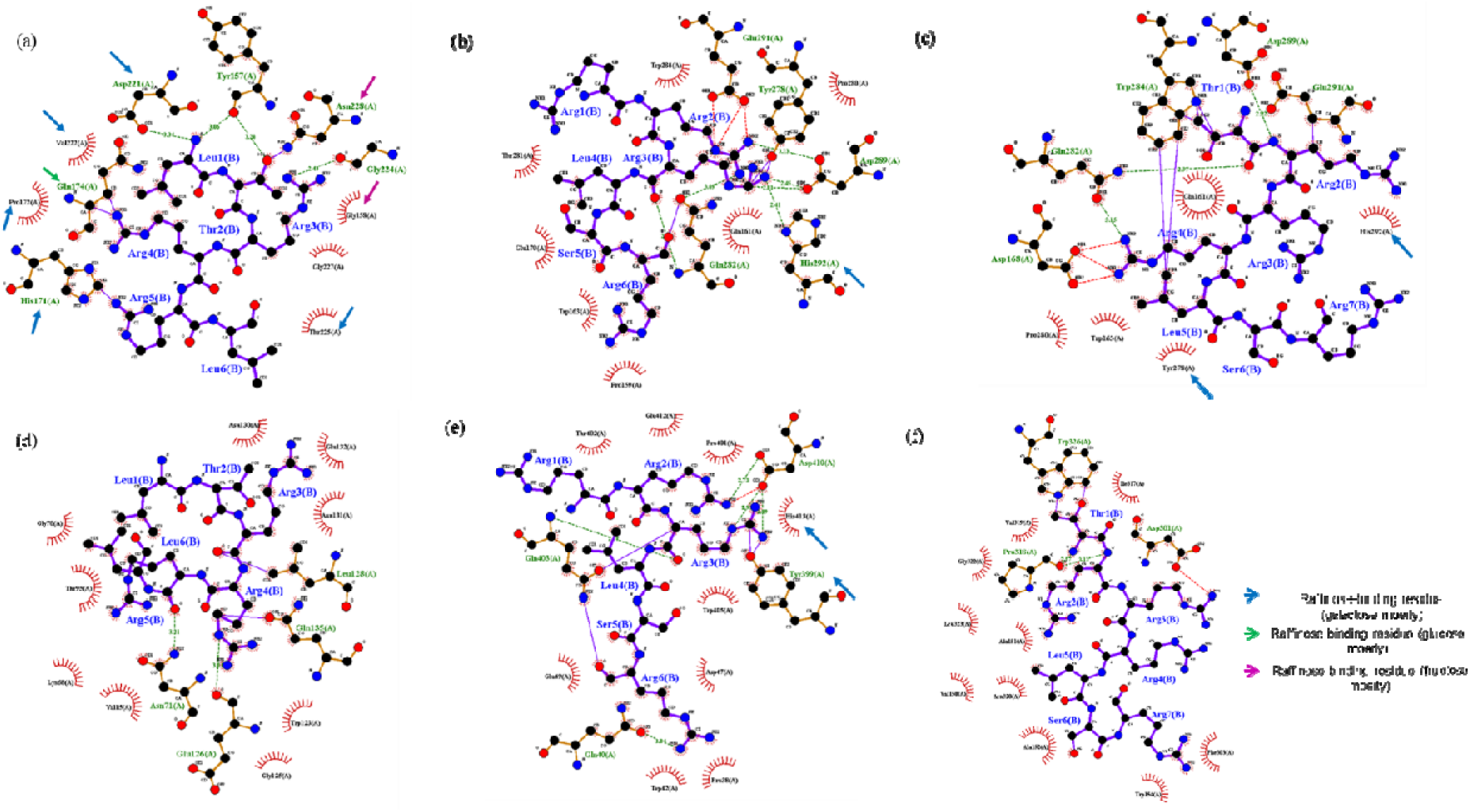
2D interaction diagram of **(a)** amp21.4 **(b)** amp21.5 **(c)** amp21.6 with LecA protein on the basis of most negative MMGBSA model. 2D interaction diagram of **(d)** amp21.4 **(e)** amp21.5 **(f)** amp21.6 with LecA protein on the basis of most negative docking score model.

**Figure 18.**
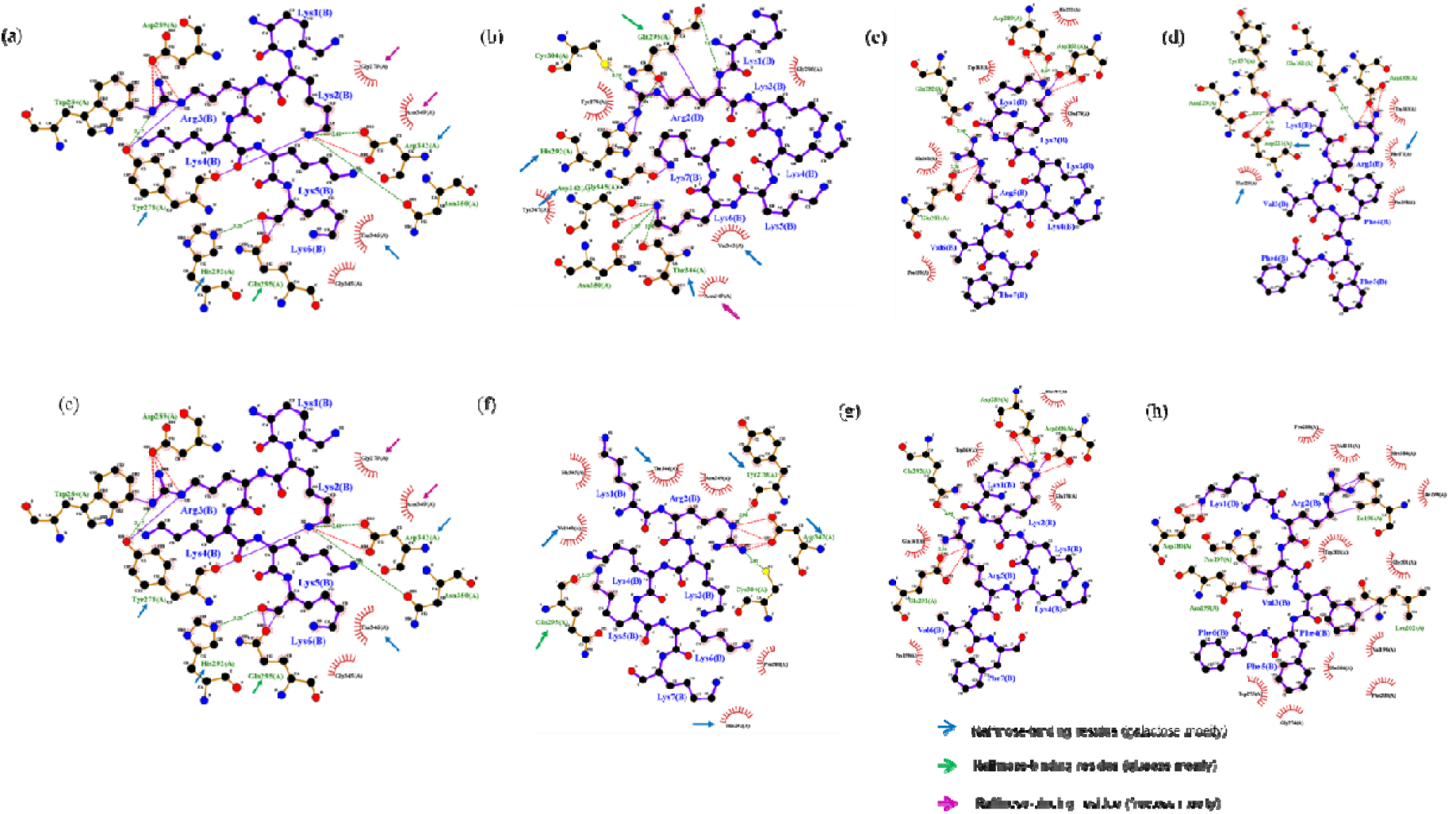
2D interaction diagram of **(a)** amp24.2 **(b)** amp24.3 **(c)** amp24.5 and **(d)** amp24.6 with LecA protein on the basis of most negative MMGBSA model. 2D interaction diagram of **(e)** amp24.2 **(f)** amp24.3 **(g)** amp24.5 and **(h)** amp21.6 with LecA protein on the basis of most negative docking score model.

#### 3.15.5. Assessment of the conformational stability and binding behavior of top-ranked USP–LecA complexes through MD simulation analysis

For the MD Simulation analysis, we have selected amp21.4, amp21.5 and amp21.6 as ultra-short derivatives of amp21. And for ultra-short derivatives of amp24, we have selected amp24.2, amp24.3, and amp24.6 for the MD simulation analysis using GROMACS.

##### 3.15.5.1 Analyzing the structural stability of USP–LecA complexes

The RMSD plots for all complexes formed between USPs of amp21 and amp24 with LecA protein are shown in Figure 19. During the 200 ns simulation, the amp21.4-LecA complex displayed noticeably lower RMSD values at several points compared to the unbound protein, indicating increased stability in those trajectory segments. The RMSD profile for the amp24.3-LecA complex was exceptionally stable; the curve stayed nearly linear with only slight oscillations during the simulation, suggesting that the complex maintained a stable conformation. Comparing the RMSD plots of remaining USPs–LecA complexes with that of the LecA protein, however, revealed greater deviations and generally higher RMSD values. These fluctuations suggest that those complexes were comparatively less stable over the simulation timeframe.

**Figure 19.**
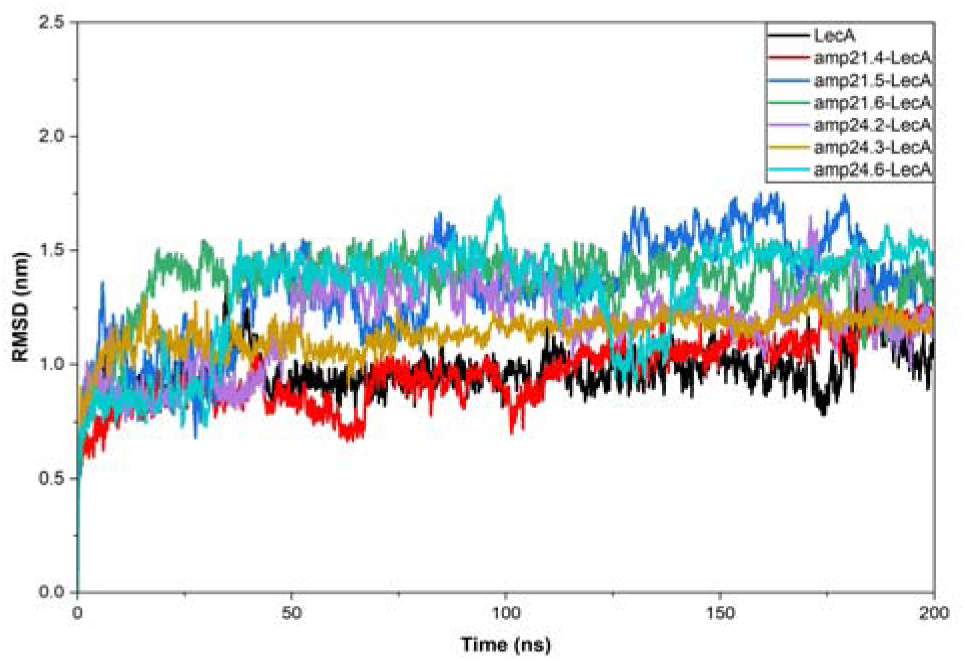
RMSD profile of USPs derived from amp21 and amp24 with LecA protein

##### 3.15.5.2. Analyzing the structural compactness of USP-LecA complexes

The radius of gyration profiles for all systems are presented in Figure 20. Across the 200 ns simulation, both the unbound LecA protein and the USP–LecA complexes showed noticeable fluctuations. At different points in the trajectory, several complexes exhibited lower Rg values than the unbound protein, while at other times their values rose above that of the apo form. Overall, the Rg values for every system, including the unbound protein, remained within a range of approximately 1.5 nm to 2.75 nm, and the curves frequently crossed over one another throughout the simulation. Because all trajectories fluctuated within a similar range and no complex showed a consistently distinct pattern, it was not possible to draw a clear conclusion about which complex maintained a more compact or stable structure based solely on the Rg analysis.

**Figure 20.**
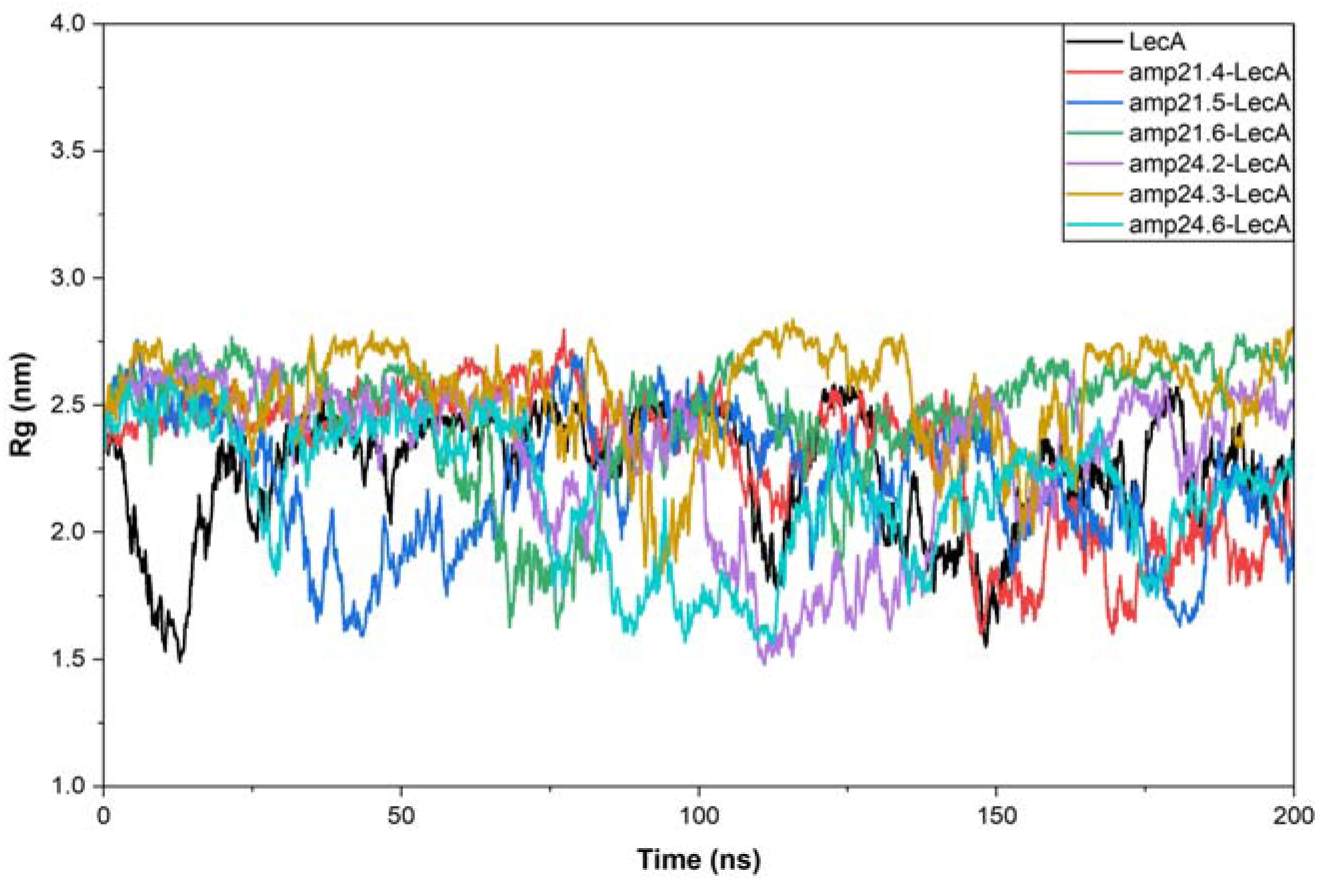
Rg profile of USPs derived from amp21 and amp24 with LecA protein

##### 3.15.5.3. Analyzing the residue-level flexibility analysis of USP–LecA Complexes

The RMSF profiles for LecA in the presence of the USPs were generally higher than those of the unbound protein, as shown in Figure 21. Among the different complexes, amp21.5-LecA and amp24.2-LecA, showed noticeably larger fluctuations at many residue positions. The amp21.4-LecA complex showed a curve that nearly overlapped with the unbound protein in the area around residues 300–350, indicating comparatively lower flexibility in that segment. Although the RMSF value of amp21.4-LecA complex was higher than that of LecA protein, the overall pattern revealed somewhat fewer fluctuations than the other peptide-bound forms. This implies that amp21.4 provides a slight stabilizing effect at specific positions and affects dynamic behaviour of LecA protein in a slightly different way than the other AMPs.

**Figure 21.**
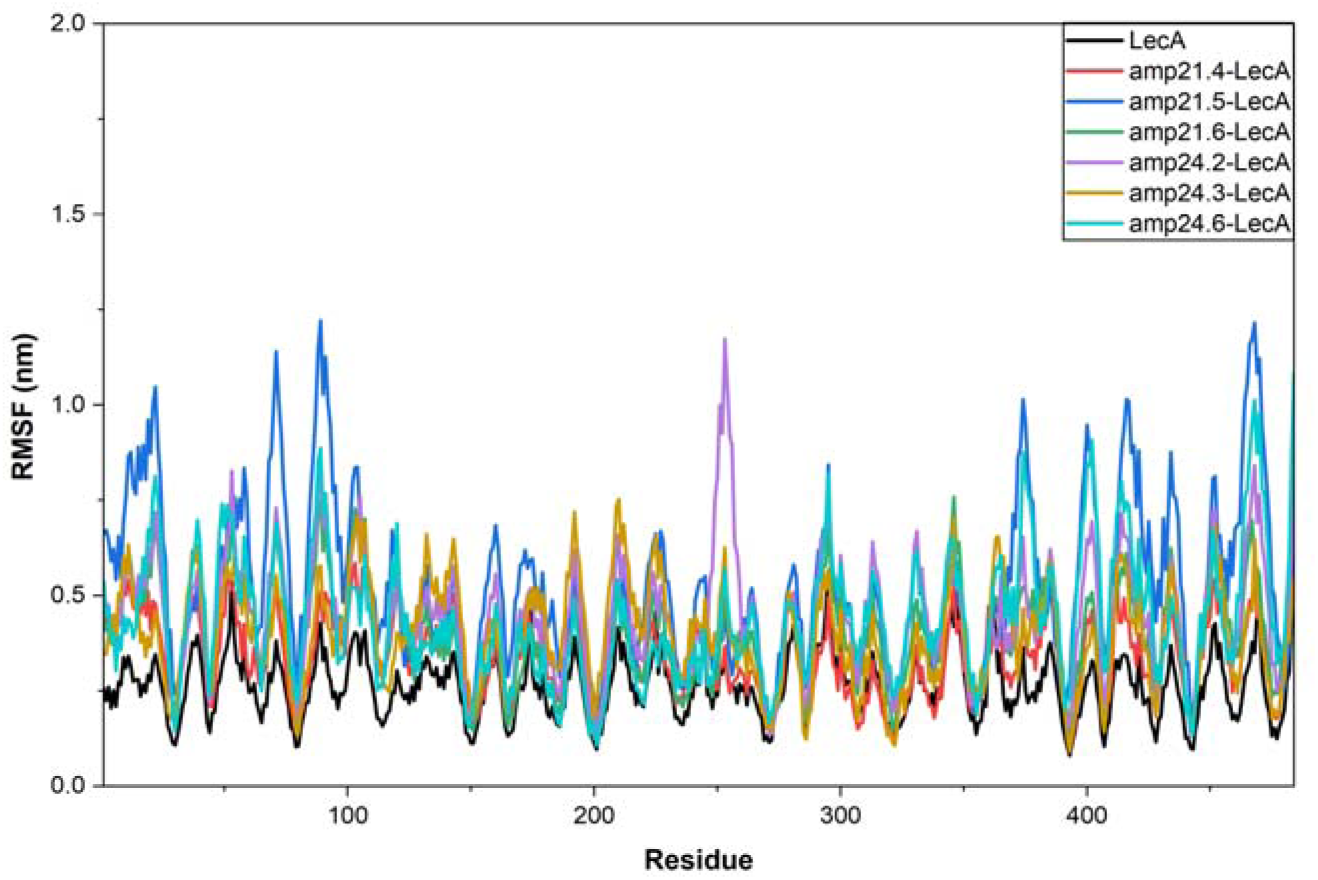
RMSF profile of USPs derived from amp21 and amp24 with LecA protein

##### 3.15.5.4. Analyzing the solvent accessibility of USP–LecA Complexes

Figure 22 displays the SASA profiles for all USP–LecA complexes as well as the unbound LecA protein. Throughout the majority of the simulation, the complexes generally showed higher SASA values than the apo protein, indicating greater surface exposure upon peptide binding. When compared to the other USPs-bound forms, the amp24.2-LecA complex displayed somewhat higher SASA values. All complexes followed a generally similar trend and continuously stayed above the unbound protein, despite some slight variations that were apparent across the trajectories. It is challenging to determine which complex performs better based solely on SASA because of this overlap in behavior.

**Figure 22.**
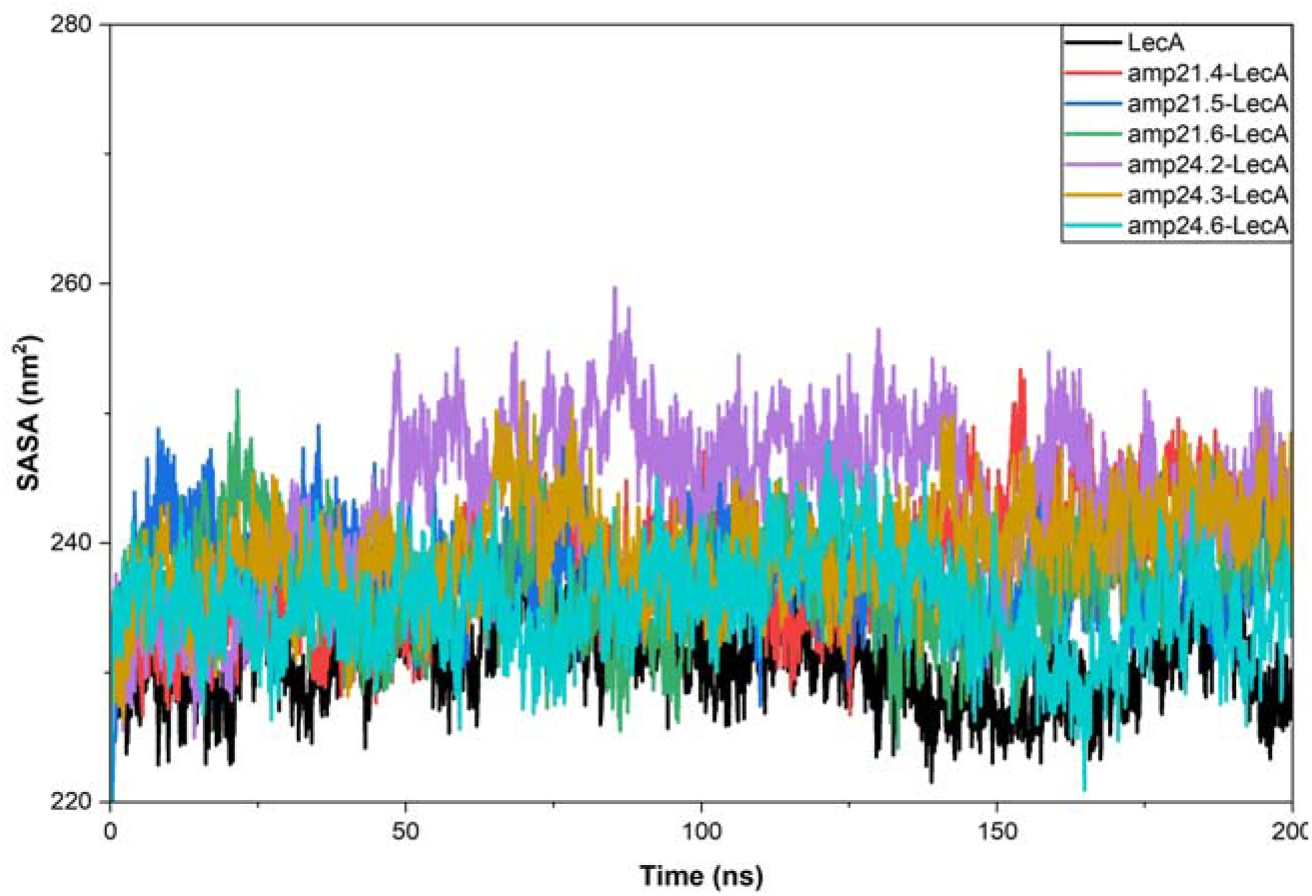
SASA profile of USPs derived from amp21 and amp24 with LecA protein

When combined, the findings from the RMSD, Rg, RMSF, and SASA analyses do not clearly show which USP, in dynamic conditions, forms a significantly more stable complex with LecA. Because the different complexes behaved in much the same way across all the parameters, it was difficult to single out any one peptide as clearly superior. The only slight pattern that emerged was in the RMSD and RMSF profiles, where amp21.4 appeared a bit more stable than the others. Even so, this difference is small and not strong enough to draw a firm conclusion from.

We then used FEL analysis, which provides a more thorough view of the conformational stability of the complexes, to confirm the validity of our results and find USPs with truly stable binding behavior.

##### 3.15.5.5. Analyzing the free energy landscape of USP–LecA Complexes

The FEL plots shown in Figure 23 give a clearer picture of how the LecA–peptide complexes behave in terms of stability. Among the amp21-derived peptides, amp21.4 stood out the most. Its PC1 and PC2 values were confined to a relatively low value of range, and its Gibbs free-energy levels were noticeably lower than those of the unbound LecA protein. When looking at the top-view of the landscape, a large portion of the conformations left the initial state of energy and gathered in the low-energy (blue) zone, suggesting that the structure settled into a stable state when this peptide was present.

**Figure 23.**
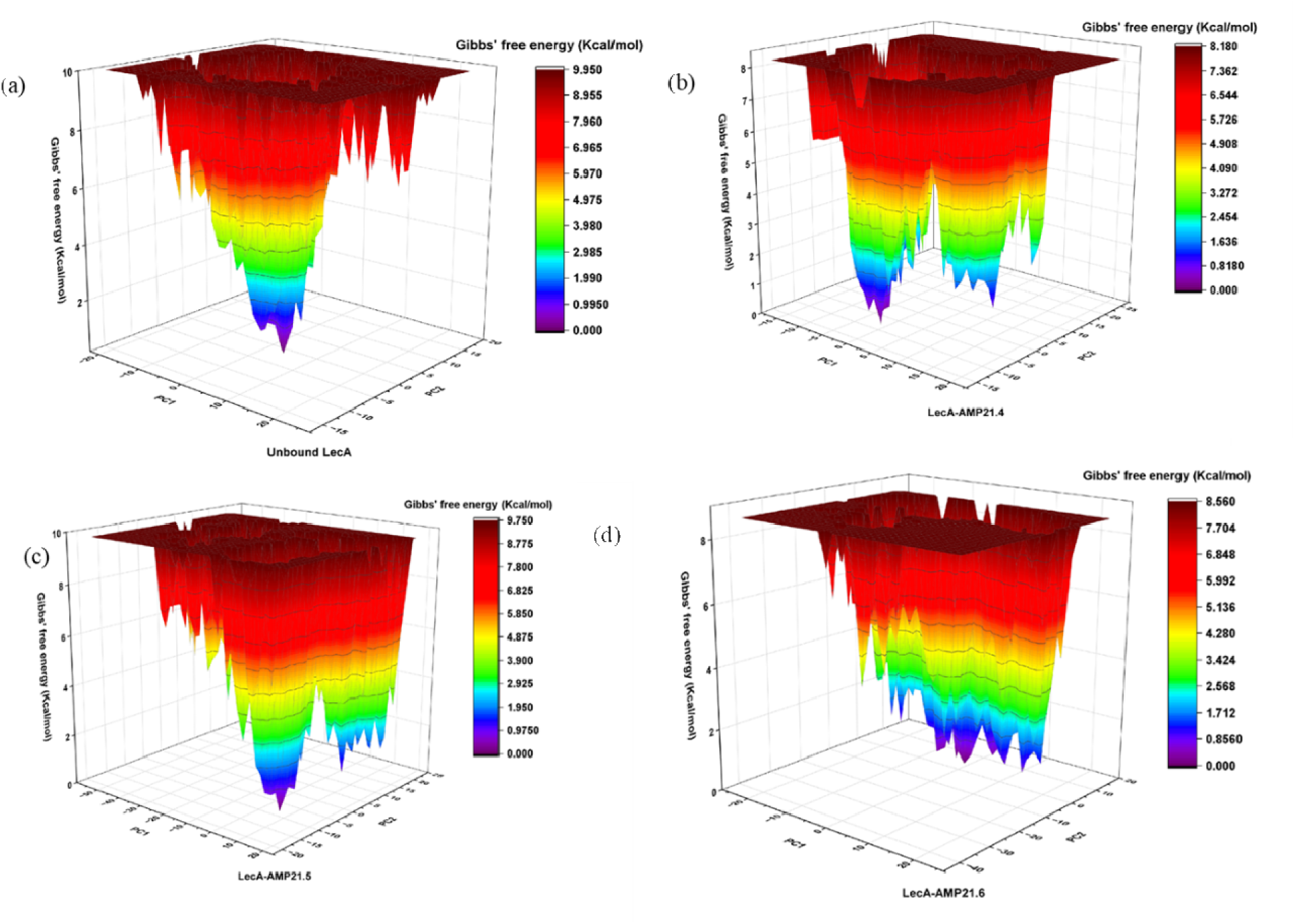
FEL plot of **(a)** unbound LecA protein **(b)** amp21.4-LecA **(c)** amp21.5-LecA and **(d)** amp21.6-LecA complex

The pattern for amp21.5 was somewhat different. Although many of its structures did reach the lower-energy region, the spread along PC1 was much broader compared with the apo protein. This wider distribution points to the protein adopting more scattered conformations when bound to amp21.5. As for amp21.6, its energy values were lower than those of the unbound protein, and its PC1 and PC2 values were within a moderate range. However, the top-view plot made it clear that many of the conformations clustered around the higher-energy region and only a few managed to move toward the lowest energy zone. This suggests that LecA does not gain much structural stability when amp21.6 is bound.

Taking all of this together, amp21.4 was the only derivative in the amp21 group that consistently maintained a compact structure and supported a stable, low-energy configuration for LecA.

A similar assessment was carried out for the USPs derived from amp24 (Figure 24). The peptide amp24.2 produced a clear shift toward more stable conformations. Its PC1 and PC2 values fell into a well-defined, narrow band, and most of its conformations clustered in the deeper energy basin and left the initial state in the top-view plot. The Gibbs free-energy range was also lower than that of the unbound protein, which strongly points to improved stability and tighter packing of LecA when amp24.2 is bound.

**Figure 24.**
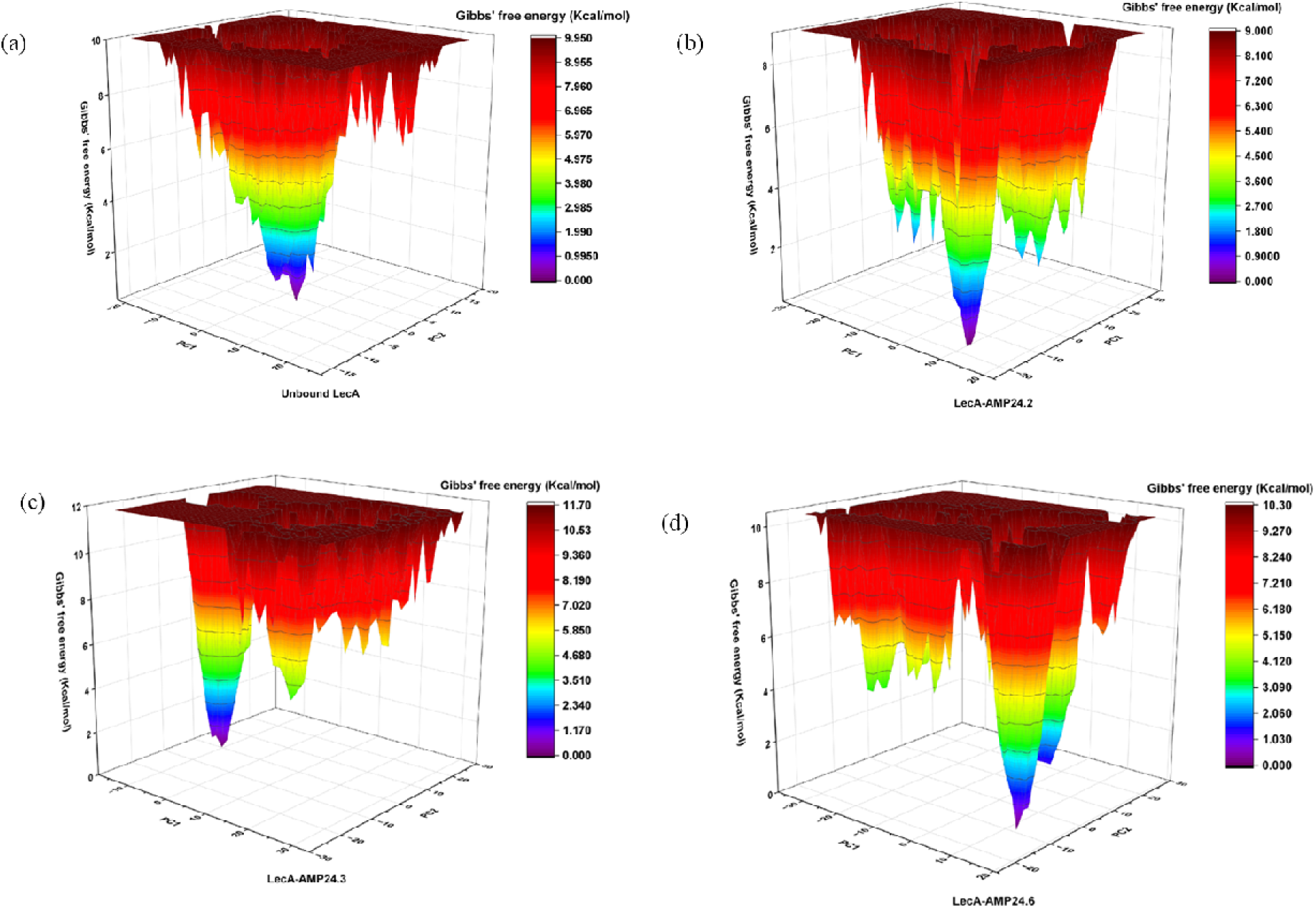
FEL plot of **(a)** unbound LecA protein **(b)** amp24.2-LecA **(c)** amp24.3-LecA and **(d)** amp24.6-LecA complex

On the other hand, amp24.3 showed a wider PC2 distribution than the apo form, indicating more spread-out conformations. Only a small portion of its structures moved into the low-energy region, and the overall Gibbs energy range remained higher, suggesting that this peptide does not stabilize the protein to any meaningful extent. A comparable trend appeared in the amp24.6 complex, where the PC values were more dispersed and the energy values were higher. Although a few conformations drifted away from the high-energy zone, very few reached the most stable energy basin, indicating a limited stabilizing influence.

Considering all these observations, amp21.4 (among the ultra-short derivatives of amp21) and amp24.2 (among the ultra-short derivatives of amp24) were the only USPs that consistently encouraged LecA to adopt stable, low-energy conformations.

#### 3.4.16. *In vitro* validation of USPs

amp21.4 and amp24.2 were found to be the potent USPs on binding to LecA protein through *in silico* studies and were therefore subjected to wet lab validation.

##### 3.4.16.1. Analyzing the hemocompatibility of USPs

Before assessing the antibiofilm efficacy of amp21.4 and amp24.2, their hemocompatibility was determined in the concentration range of 12.5-100 µg mL^-1^ using our previously standardized protocol [Singh et al., 2022]. It was found that both amp21.4 and amp24.2 showed comparable hemocompatibility with that of the parent AMPs i.e. amp21 and amp24 (Figure 25.). However, as USPs possess much smaller size as compared to the parent AMPs, it is conceivable that they are likely to be more biocompatible with the parent AMPs.

**Figure 25.**
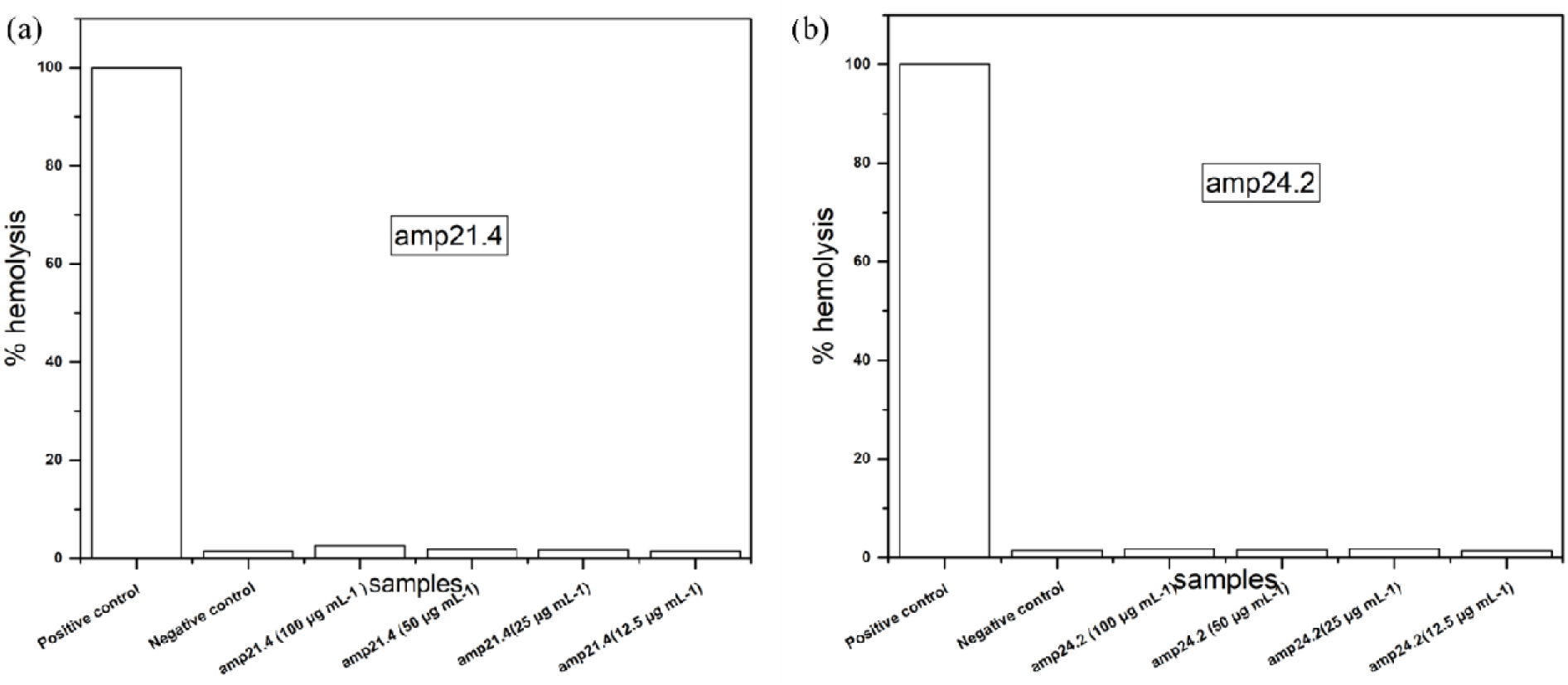
Hemolysis assay of amp21.4 and amp24.2

##### 3.4.16.2. Analyzing the antibiofilm efficacy of USPs (amp21.4 and amp24.2)

The antibiofilm efficacy of the USPs (amp21.4 and amp24.2) were assessed quantitatively against *P. aeruginosa* through both biofilm inhibition and disruption studies following our standard protocol. Since USPs are extremely small in size, they are likely to be less cytotoxic at higher concentrations. Therefore, we investigated the biofilm inhibition and biofilm disruption potency of both amp21.4 and amp24.2 at 250 µg mL^-1^ and compared their efficacy with the parent AMPs at same concentration.

Biofilm inhibition assays showed amp21.4 and amp24.2 reduced the biofilm mass to 41.29% and 41.38% respectively of the untreated control that was considered to be 100%respectively (Figure 26a). On the other hand, amp21 and amp24 reduced the biofilm mass to 44.22% and 48.04% of control respectively (Figure 26a). These findings indicate that when compared to the parent AMPs, both the USPs i.e. amp21.4 and amp24.2 exhibited better biofilm inhibition activity, with the improvement being more pronounced for amp24.2.

**Figure 26.**
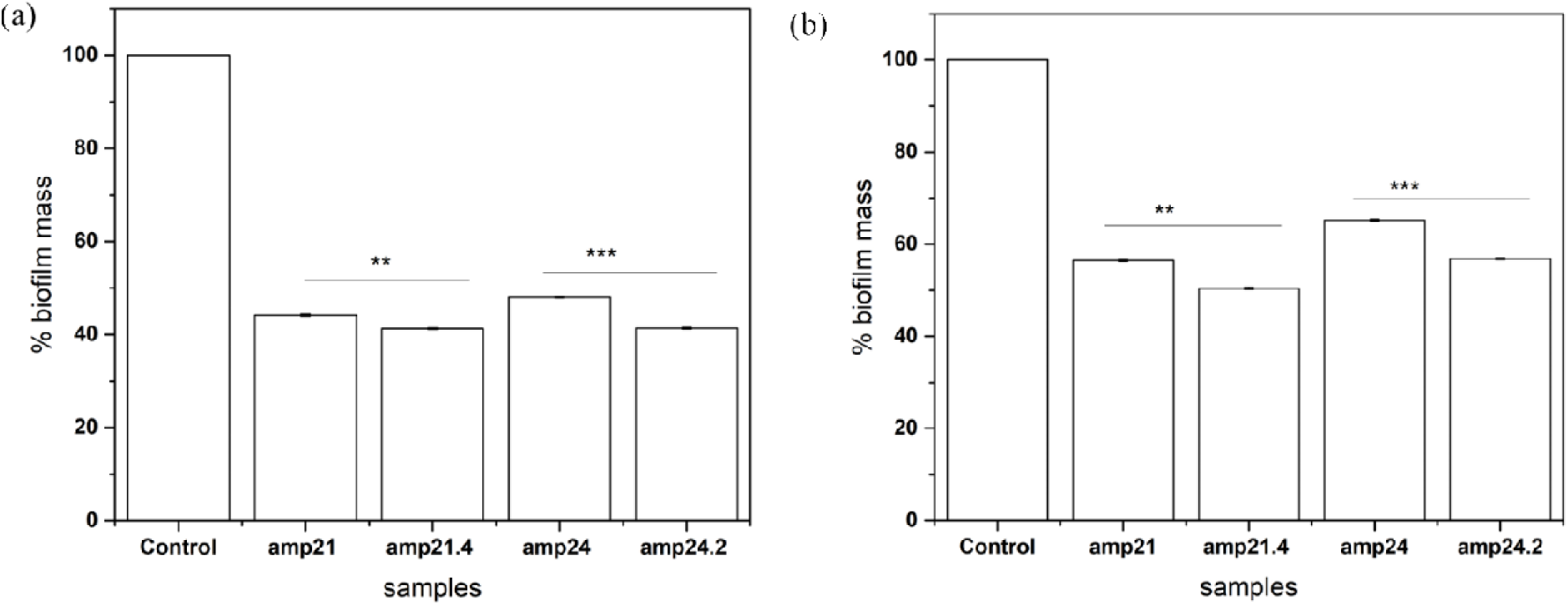
**(a)** Comparative study of *P. aeruginosa* biofilm inhibition by amp21.4 and amp24.2 with their parent AMPs at 250 µg mL^-1^ **(b)** Comparative study of *P. aeruginosa* biofilm disruption by amp21.4 and amp24.2 along with their parent AMPs at 250 µg mL^-1^

In the biofilm disruption experiment, treatment with amp21.4 and amp24.2 reduced the biofilm mass to 50.41% and 56.82% respectively of the untreated control (Figure 26b). In contrast, the corresponding parent AMPs reduced the biofilm mass to 56.5% and 65.23% respectively (Figure 5.26b). The above findings demonstrated that the USPs i.e. amp21.4 and amp24.2 were more effective in disrupting pre-formed biofilms as compared to the parent AMPs. amp24.2 was found to show a more noticeable enhancement in disrupting the preformed biofilms.

## 4. Conclusion and future perspective

AMPs from the human gut microbiome were computationally identified as potent inhibitors of lectins that are key targets in biofilm formation in *Pseudomonas aeruginosa*. *In vitro* studies showed that the lead AMPs filtered through *in-silico* studies showed strong antibiofilm activity by stimulating the migration of bacterial cells from the sessile biofilm stage to the mobile planktonic stage. Apart from disrupting the biofilm EPS matrix, the above AMPs exhibited prominent antibacterial activity by compromising the integrity of the bacterial cell membrane. For practical applications, the efficacy of the above lead AMPs can also be investigated in an animal model.

## Supplementary information

All relevant supplementary information has been incorporated in Supplementary file 1

## Acknowledgment

AA would like to thank ICMR, Govt. of India, for his fellowship. AAA and SC would like to thank the MoE, Govt. of India, for their fellowship. SA would like to thank CSIR, Govt. of India, for his fellowship. We would like to thank IIIT Allahabad for providing the research and the Central Computation Facility (CCF). We would like to thank the Electron Microscopy Unit of CDRI Lucknow for providing the SEM imaging facility.

## Declaration of generative AI and AI-assisted technologies in the writing process

During the preparation of this work, the author(s) used ChatGPT to assist with language refinement and improve clarity. After using this tool/service, the author(s) reviewed and edited the content as needed and take(s) full responsibility for the content of the publication.

