## Supplementary material for "Identification of Human Gut Microbiome–Derived Peptides Targeting Biofilm-Specific Lectin Proteins of *Pseudomonas aeruginosa*": main.py

```
from Bio import SeqIO

def extract_sequences_by_keywords(input_fasta, output_fasta, keywords_file):
    # Read the list of keywords from the file
    with open(keywords_file, 'r') as file:
        keywords = set(line.strip() for line in file)

    # Parse the input FASTA file and write the matching sequences to the output file
    with open(output_fasta, 'w') as out_file:
        for record in SeqIO.parse(input_fasta, 'fasta'):
            if any(keyword in record.description for keyword in keywords):
                SeqIO.write(record, out_file, 'fasta')

if __name__ == "__main__":
    input_fasta = "input.fasta"   # Path to the input FASTA file
    output_fasta = "output.fasta" # Path to the output FASTA file
    keywords_file = "keywords.txt" # Path to the file containing the list of keywords

    extract_sequences_by_keywords(input_fasta, output_fasta, keywords_file)
```
