## Supplementary figures and images for "Identification of Human Gut Microbiome–Derived Peptides Targeting Biofilm-Specific Lectin Proteins of *Pseudomonas aeruginosa*"

### circular_without_bl.pdf

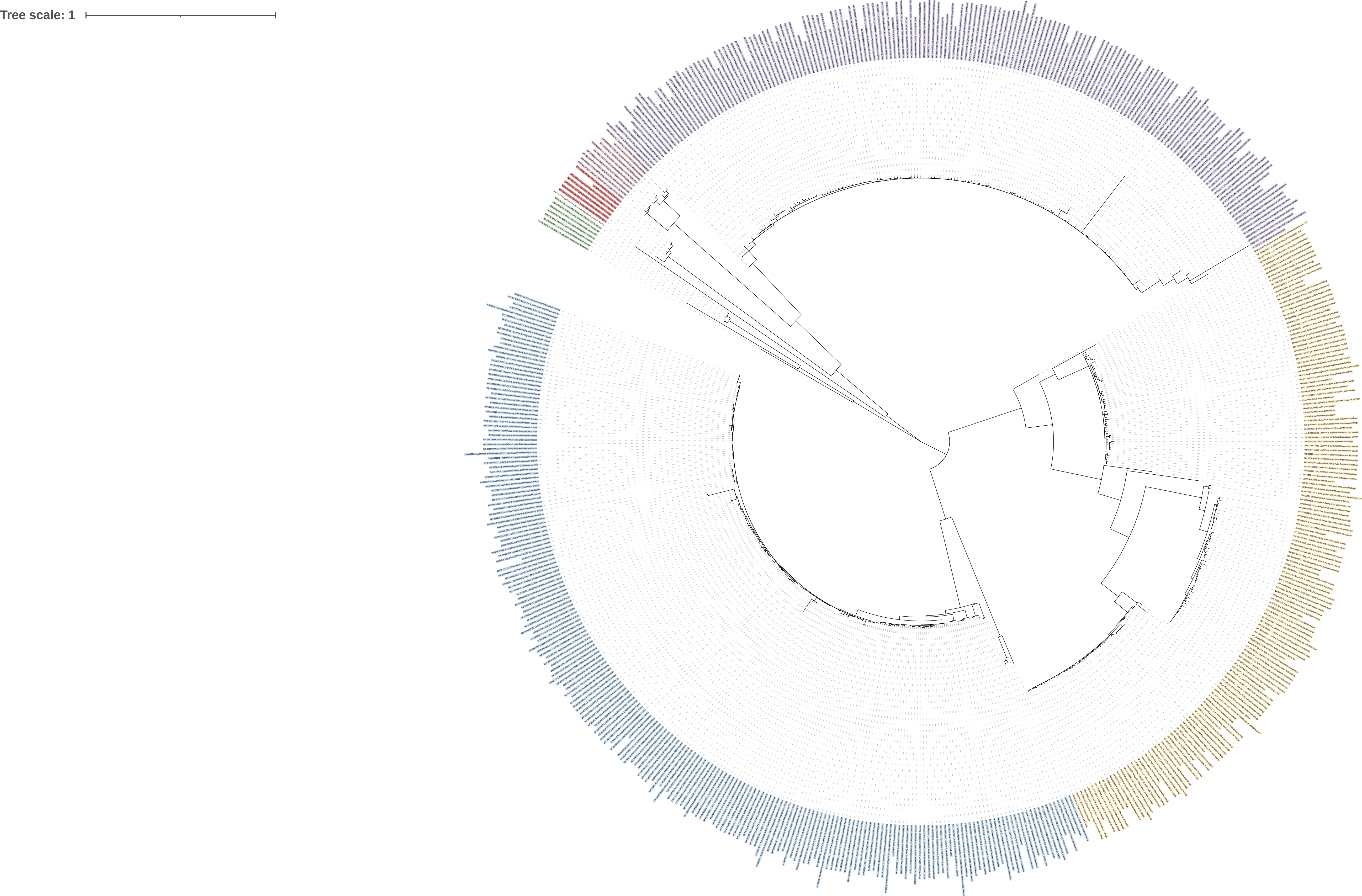

### circular_without_bl.pdf

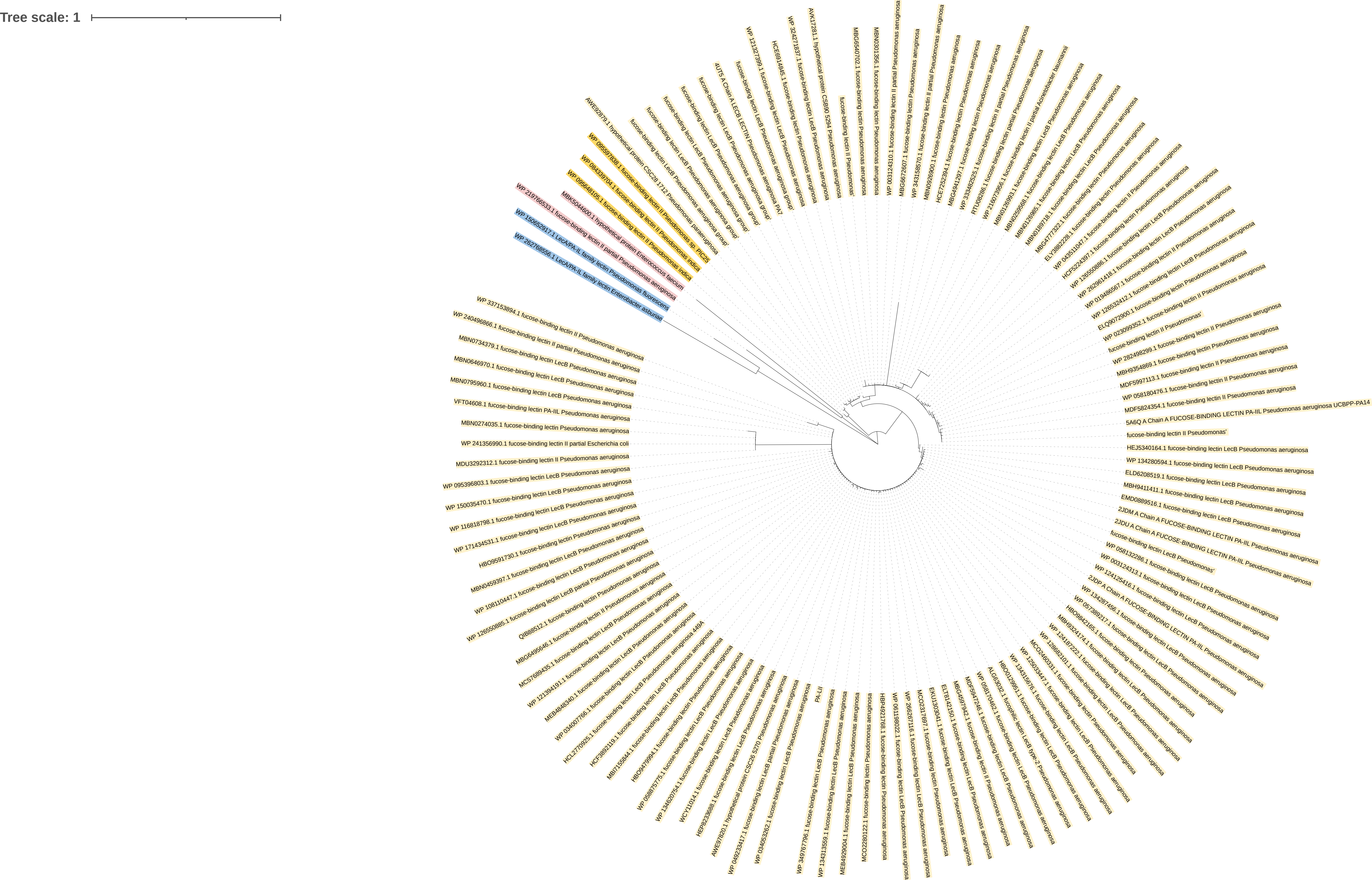

### rectangular_tree_rooted.pdf

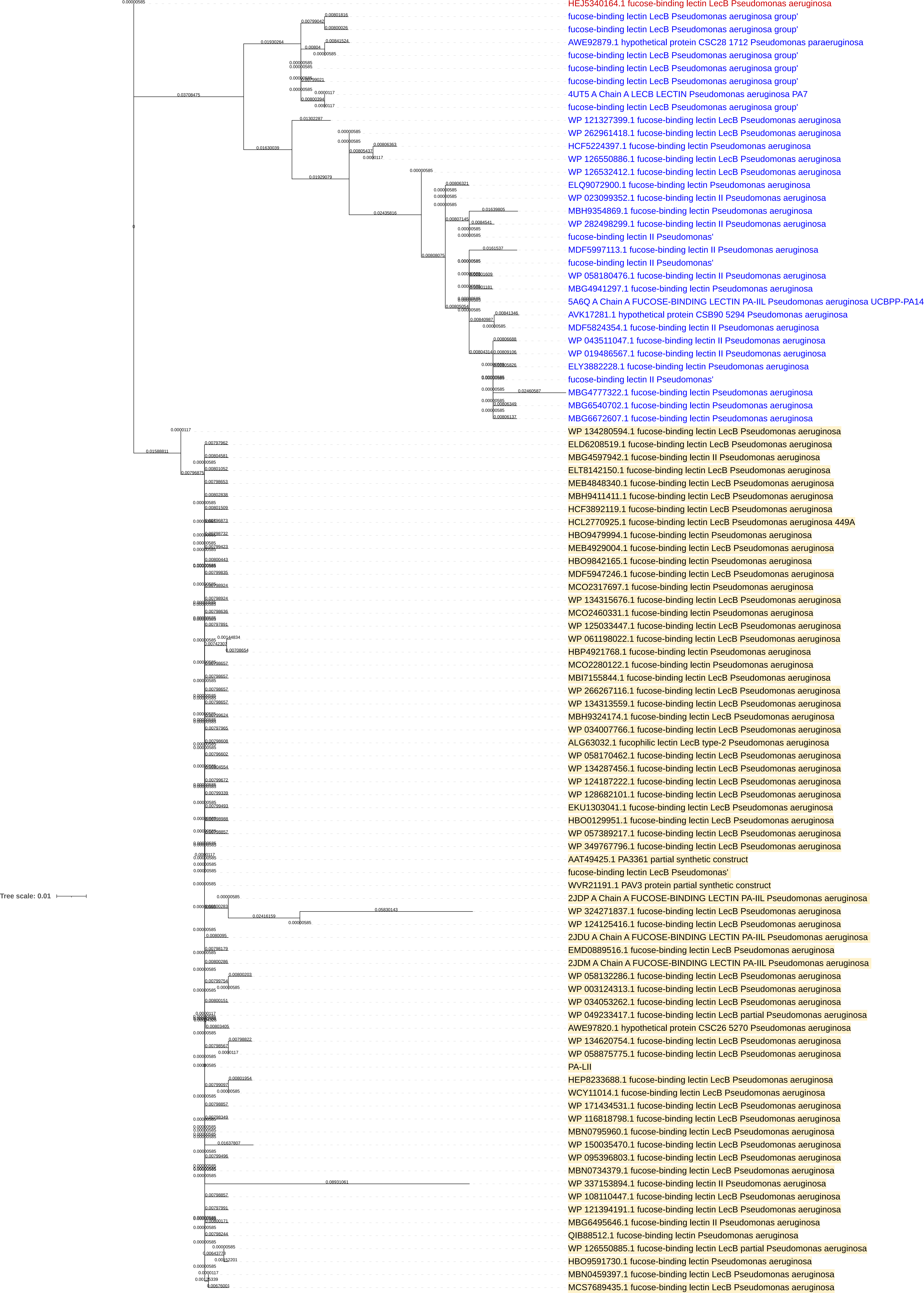

### rectangular_with_bl.pdf

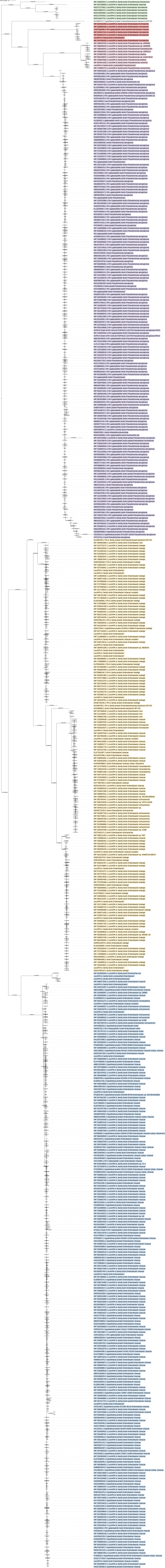

### rectt_with_bl.pdf

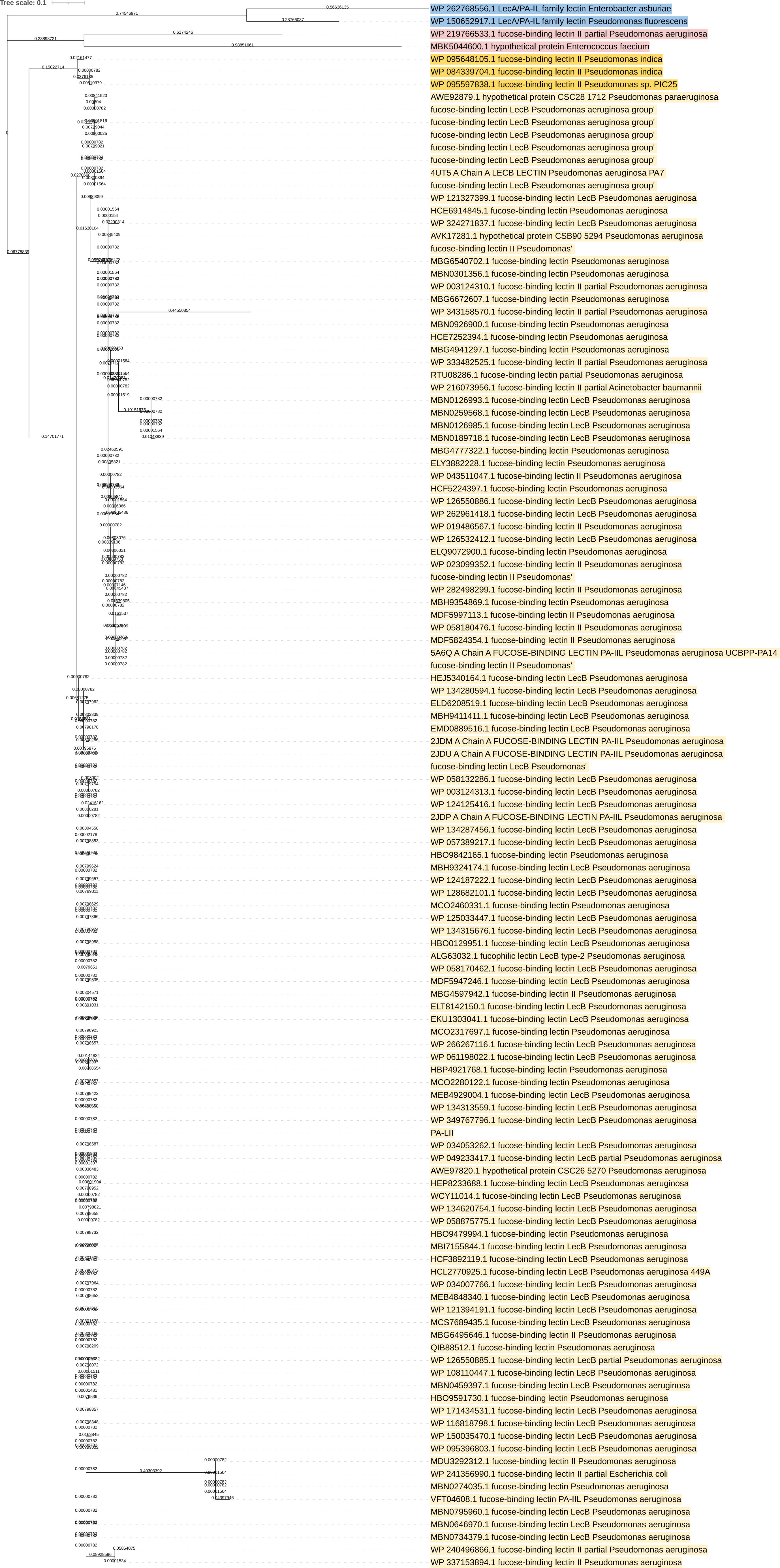

### tree_rectangular_rooted.pdf

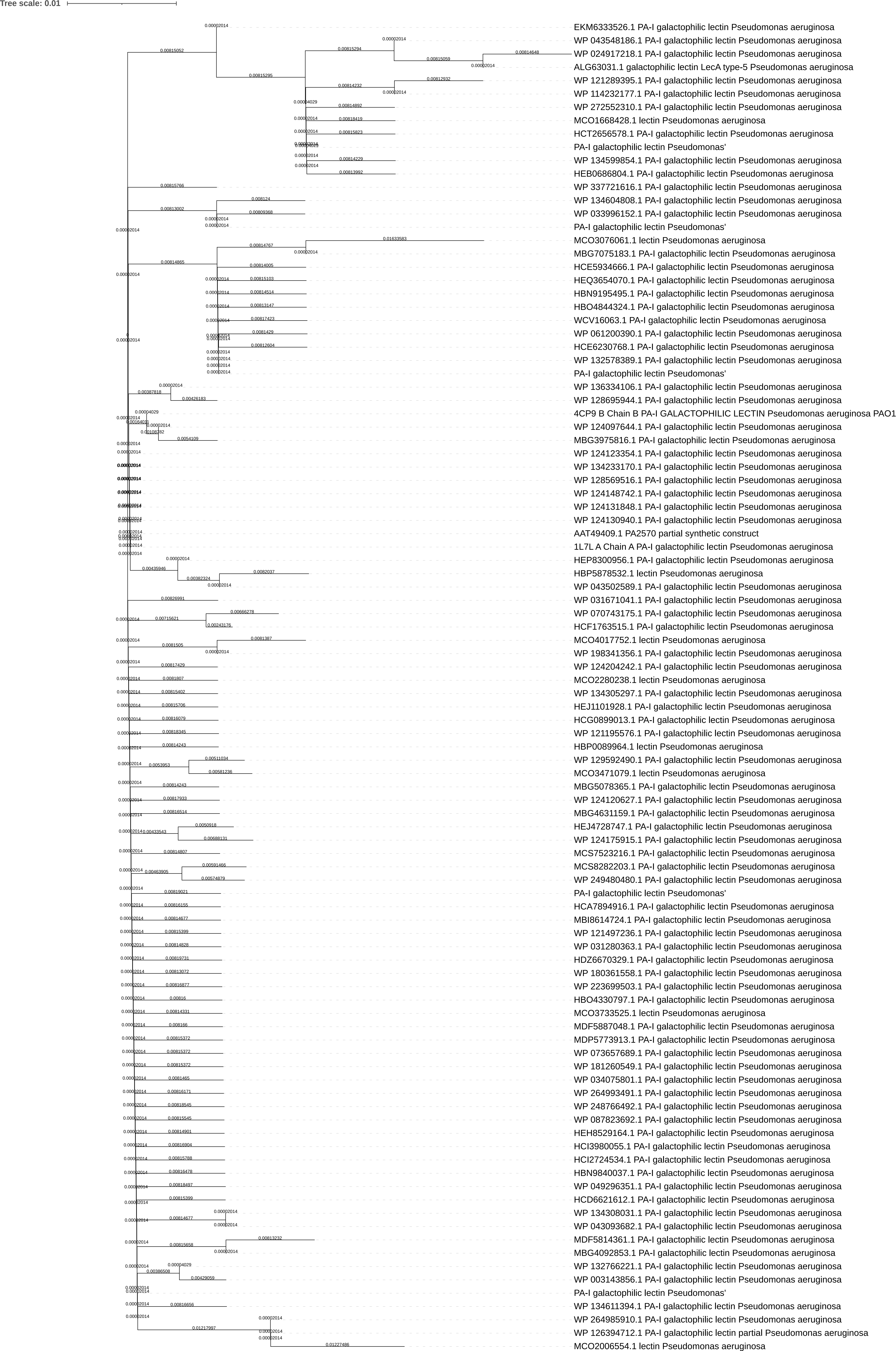
